# Genome and transcriptome analysis of the mealybug *Maconellicoccus hirsutus*: A model for genomic Imprinting

**DOI:** 10.1101/2020.05.22.110437

**Authors:** Surbhi Kohli, Parul Gulati, Jayant Maini, Shamsudheen KV, Rajesh Pandey, Vinod Scaria, Sridhar Sivasubbu, Ankita Narang, Vani Brahmachari

## Abstract

In mealybugs, transcriptional inactivation of the entire paternal genome in males, due to genomic imprinting, is closely correlated with sex determination. The sequencing, *de-novo* assembly and annotation of the mealybug, *Maconellicoccus hirsutus* genome and its comparison with *Planococcus citri* genome strengthened our gene identification. The expanded gene classes, in both genomes relate to the high pesticide and radiation resistance; the phenotypes correlating with increased gene copy number rather than the acquisition of novel genes. The complete repertoire of genes for epigenetic regulation and multiple copies of genes for the core members of polycomb and trithorax complexes and the canonical chromatin remodelling complexes are present in both the genomes. Phylogenetic analysis with *Drosophila* shows high conservation of most genes, while a few have diverged outside the functional domain. The proteins involved in mammalian X-chromosome inactivation are identified in mealybugs, thus demonstrating the evolutionary conservation of factors for facultative heterochromatization. The transcriptome analysis of adult male and female *M.hirsutus* indicates the expression of the epigenetic regulators and the differential expression of metabolic pathway genes and the genes for sexual dimorphism. The depletion of endosymbionts in males during development is reflected in the significantly lower expression of endosymbiont genes in them.

**Author summary:** The mealybug system offers a unique model for genomic imprinting and differential regulation of homologous chromosomes that pre-dates the discovery of dosage compensation of X chromosomes in female mammals. In the absence of robust genetics for mealybugs, we generated and analysed the genome and transcriptome profile as primary resources for effective exploration. The expanded gene classes in the mealybugs relate to their unique biology; the expansion of pesticide genes, trehalose transporter, SETMAR and retrotransposons correlate with pesticide, desiccation and radiation resistance, respectively. The similarity in the genomic profile of two species of mealybugs strengthens our gene prediction. All the known epigenetic modifiers and proteins of the primary complexes like the PRC1,2 and the trithorax are conserved in mealybugs, so also the homologues of mammalian proteins involved in X chromosome inactivation. The high copy number of genes for many partners in these complexes could facilitate the inactivation of a large part of the genome and raise the possibility of formation of additional non-canonical complexes for sex specific chromosome inactivation. In adult males and females, the status of epigenetic regulation is likely to be in a maintenance state; therefore, it is of interest to analyze the expression of epigenetic regulators during development.

## Introduction

The mealybugs (Hemiptera:Pseudococcidae), such as *Maconellicoccus hirsutus* (Mhir)and *Planococcus citri* (Pcit), commonly feed on plant sap. They are considered as invasive species having a wide host-range and are spread in all parts of the world. *M. hirsutus* and *P.citri* commonly reproduce sexually, though parthenogenesis is reported in *M. hirsutus*[1, 2]. The life cycle is completed in around 29 days at 27^0^C. The insect predators are most often used for the control of these invasions.

The adult mealybugs are sexually dimorphic with males being small & winged and the females being much larger, wingless & sedentary. The immature males and females (commonly known as crawlers) are morphologically similar. The males undergo four stages of metamorphosis to become adults while the female passes through only three stages along with the growth in size [3].

The chromosomal cycle of the mealybugs is a point of interest(Fig 1) as the diploid genome of mealybugs consists of five pairs of chromosomes (2n=10) and do not have any morphologically distinct sex chromosomes. However, there is sex-specific heterochromatization and transcriptional silencing of the paternally inherited chromosomes in males. Thus, genomic imprinting and the differential regulation of homologous chromosomes operate on 50% of the genome. The heterochromatization and transcriptional silencing of paternal chromosomes in male mealybugs is comparable to X chromosome inactivation in female mammals [4–7]. Unlike X chromosome inactivation, paternal genome inactivation is non-random in the mealybugs. X chromosome inactivation in mammals and the paternal genome inactivation in mealybugs both result in physiological haploidy and differential regulation of homologous chromosomes within the same nucleus. The attributes of genomic imprinting in the mealybugs are described through elegant cytological and molecular studies [5, 6]. The mealybugs have holocentric chromosomes and exhibit extreme meiotic drive. During spermatogenesis in males, it is only the maternally inherited chromosomes, which contribute to the active sperms, while the paternally inherited chromosomes undergo heteropycnosis, leading to their disintegration [8]. Mealybugs are also important model organisms to study responses to high doses of ionizing radiation as they can tolerate radiation doses ∼1100Gy [9].

**Fig 1.**
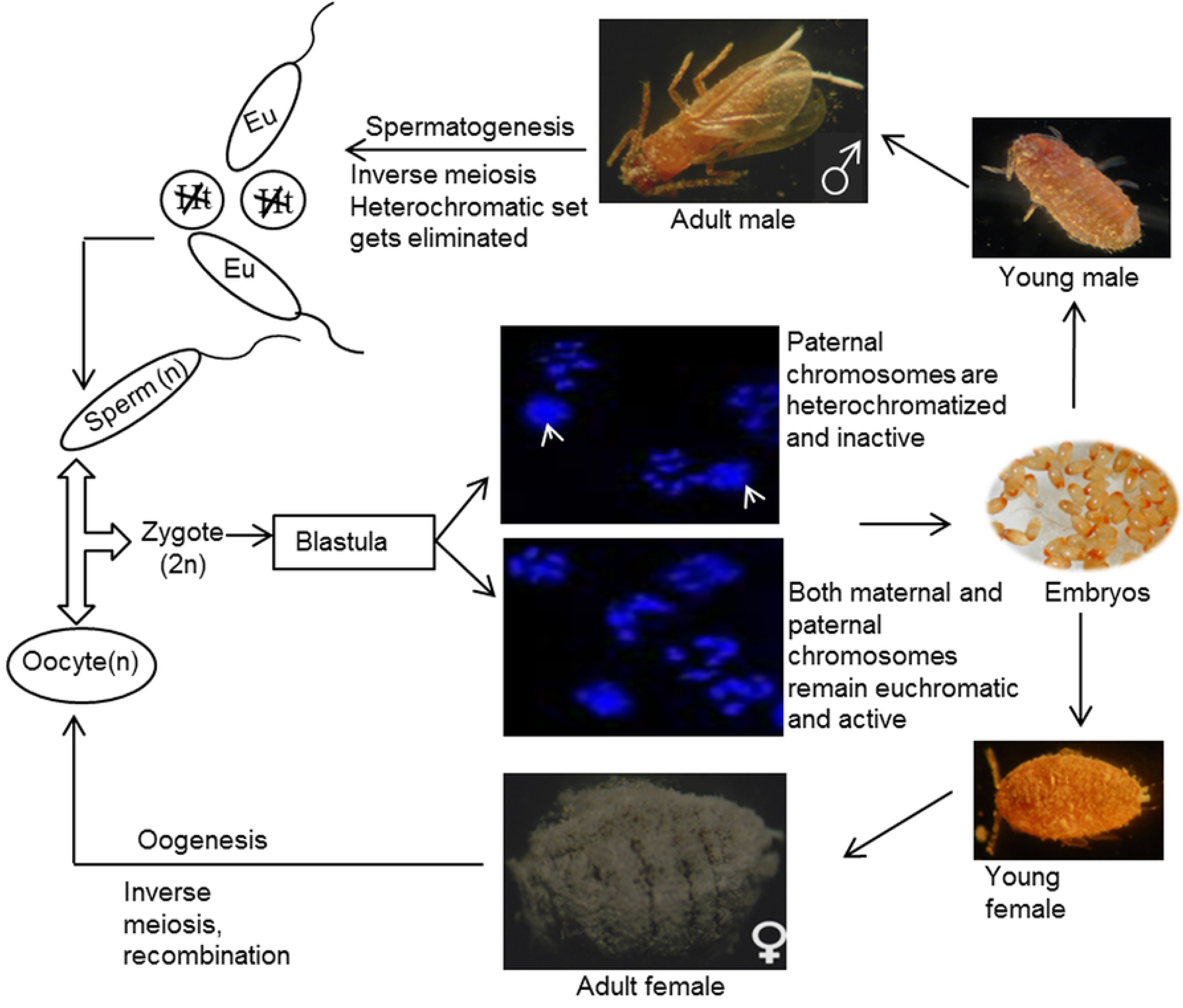
**Life cycle of the mealybug Maconellicoccus hirsutus.** The developmental stages are similar between different species of mealybugs. The heterochromatin in males is indicated (white arrow). The inactive and condensed paternal genome does not contribute to mature sperms, Eu-maternal euchromatin, Ht-paternal heterochromatin [7].

The various molecular features relating to genomic imprinting and epigenetics in mealybugs have been investigated by different groups. Both DNA methylation and post-translational modification of histones are predicted to be involved in imprinting in this system [10–13]. In *P. lilacinus*, DNA methylation was detected in CpA and CpT dinucleotides in addition to CpG [14]. The role of post-translational modification of histones in heterochromatin in paternal nuclei is highlighted in multiple studies [13, 15]. The presence of a male specific chromatin organization designated as Nuclease Resistant Chromatin (NRC) is demonstrated in two species of mealybugs, *Planococcus lilacinus* and *Maconellicoccus hirsutus*[13, 16]. NRC is predicted to include potential centers of inactivation for heterochromatin formation in male mealybugs [5, 16].

All mealybug species live in symbiosis with the β-proteobacterium *Tremblaya* and this β-proteobacterium harbours additional γ-proteobacterial species (like *Moranella*nPcit*; Doolittlea endobia* in Mhir) [17, 18]. The detailed genomic analysis of the microbiome of mealybugs has revealed extensive metabolic cooperation between mealybugs and their endosymbionts [17, 18].In the work described here, we generated the primary data by sequencing and annotating the mealybug genome with a focus on genes involved in epigenetic regulation and genomic imprinting. This will complement the molecular and the immuno-microscopic studies that are pursued, in absence of a robust genetic analysis in this system. We sequenced and annotated the genome of the pink mealybug, *Maconellicoccus hirsutus* (Mhir). We annotated the sequence of *Planococcus citri* (Pcit) genome sequence given by Husnik and McCutcheon [18]. We annotated predicted genes using BLASTp and identified functional domains using InterPro and compared the annotation of both methods.This enableda better prediction of function of genes which BLASTp could not annotate. We analyzed specific classes of genes related to different aspects of the biology of mealybugs, mainly epigenetic regulation.

We analysed the genome for horizontal gene transfers (HGTs) and expansion and contraction of gene classes. Along with the HGTs identified earlier, we found novel HGTs coding for antioxidant enzymes, protease inhibitors, bacterial toxins and carbohydrate metabolism proteins. The pesticide resistance gene classes are identified as one of the expanded classes in the genome. We performed comparative analysis of selected gene classes between Mhir, Pcit*, A.pisum* (Apis)*, C. lectularius* (Clec) *and D.melanogaster* (Dmel), which showed that the epigenetic machinery in the mealybug is complete, including the writers, readers and the erasers. A comparative transcriptome analysis indicates that these genes are expressed in both adult males and females, while some are differentially expressed.

## Results and discussion

### Genome assembly, evaluation and validation

The de novo assembly of mealybug genomewas carried out using a hybrid approach using the MaSuRCA pipeline as described under methods. The length distribution of PacBio reads are provided in Table 1. Error correction of low coverage PacBio reads (5.48x) was done using high coverage Illumina data (56.3X) in a sensitive mode. Further, error corrected PacBio reads were merged with Illumina super-reads using Celera assembler. There were 214,820 error corrected reads which have length ≥500bp that were assembled with Illumina super-reads to generate the final assembly.There were 7747 scaffolds that contributed to 168.28 Mb assembly with N50 of ∼57 Kb. Other parameters of assembly statistics are provided in Table 1. This estimate is very close to the predicted genome size of ∼163 Mb for *M. hirsutus* genome [18]. In addition, 27,885 degenerate (degen) contigs were also added to the main assembly to ensure the completeness of assembly in terms of genes. The degen contigs were not part of scaffolds due to low coverage. We mapped back Illumina reads to degen contigs and found that ∼95% of degen contigs mapped to the Illumina reads which contributed ∼21.8 Mb to the assembly. Thus, confirming that degen contigs are derived from Mhir genome.

**Table 1:** Summary of the genome sequencing data.

| Platform | Read length |  |
| --- | --- | --- |
|  | Raw reads | Filtered reads |
| **IlluminaHiSeq (Paired-End)** | 100149316 (101bp) | 84930794 (50-101bp) |
| **PacBio** | 1,60,775 (50-544,91bp) | 2,14,820(500-39578bp) |
| **Ion Torrent (Single-End)** | 14,913,519 (8-745bp) | 10,814,178 |
| **N50** | 57,095bp or ~57 Kb |  |
| **Assembly size\* (bp)** | 189.24 Mb |  |
| **No.of scaffolds** | 28,882 |  |
| **Largest scaffold** | 523,004 bp or 0.52 Mb |  |
| **BUSCOs** | C:72% [D:5.6%], F:17%, M:10%, n:2675 |  |
| **Predicted unique genes** | 21,623 |  |
| **Repeats Masked** | 19.96% |  |

The details of the sequence data pre- and post-filtering for quality are given against each platform used. The number in parenthesis indicates the size range of sequence reads. Under the BUSCOs, C- Complete single-copy BUSCOs, D- Complete duplicated BUSCOs, F- Fragmented BUSCOs, M-Missing BUSCOs, n-Total BUSCO groups studied. \*Endosymbiont scaffolds are included within assembly and are tagged appropriately.

We further evaluated the completeness of assembly using different approaches. Mhir genome was also sequenced and assembled using IonTorrent (S1 Table). We aligned 18,816 Ion Torrent contigs from 10,814,178 filtered reads on the mealybug genome assembly. 96.38% of contigs aligned to the genome assembly, substantiating its completeness. The presence of single copy orthologs from phylum Arthopoda in Mhir was estimated using BUSCO. There were 72% complete BUSCOs while 10% were missing. The genome sequence can be accessed at NCBI, Genbank accession number GCA-003261595.1.

We predicted 22,723 transcripts (21,623 unique genes) in the mealybug genome using BRAKER out of which we could annotate 17,661 genes using BLASTp with NR database. After applying a filter of 30% identity and 50% query coverage to remove erroneous hits, 15,142 transcripts remained. There were 14,010 transcripts retained after removing 1,132 truncated entries. Those transcripts that did not have any domain and had coverage of less than 30% of reference sequence in BLAST were considered as truncated.

The assembly was validated using Sanger sequencing following amplification of the histone gene cluster by tiling PCR (Fig 2). A scaffold (scf7180000076114) containing all the core histone genes was identified and eight pairs of primers were designed, each spanning a region of approximately 2Kb with overlapping end sequences, covering a total length spanning 13Kb(Fig 2). The amplicons were sequenced and mapped back to the scaffold in the assembly, to confirm their organization. The gene organization was also confirmed by long PCR (Fig 2 I).

**Fig 2.**
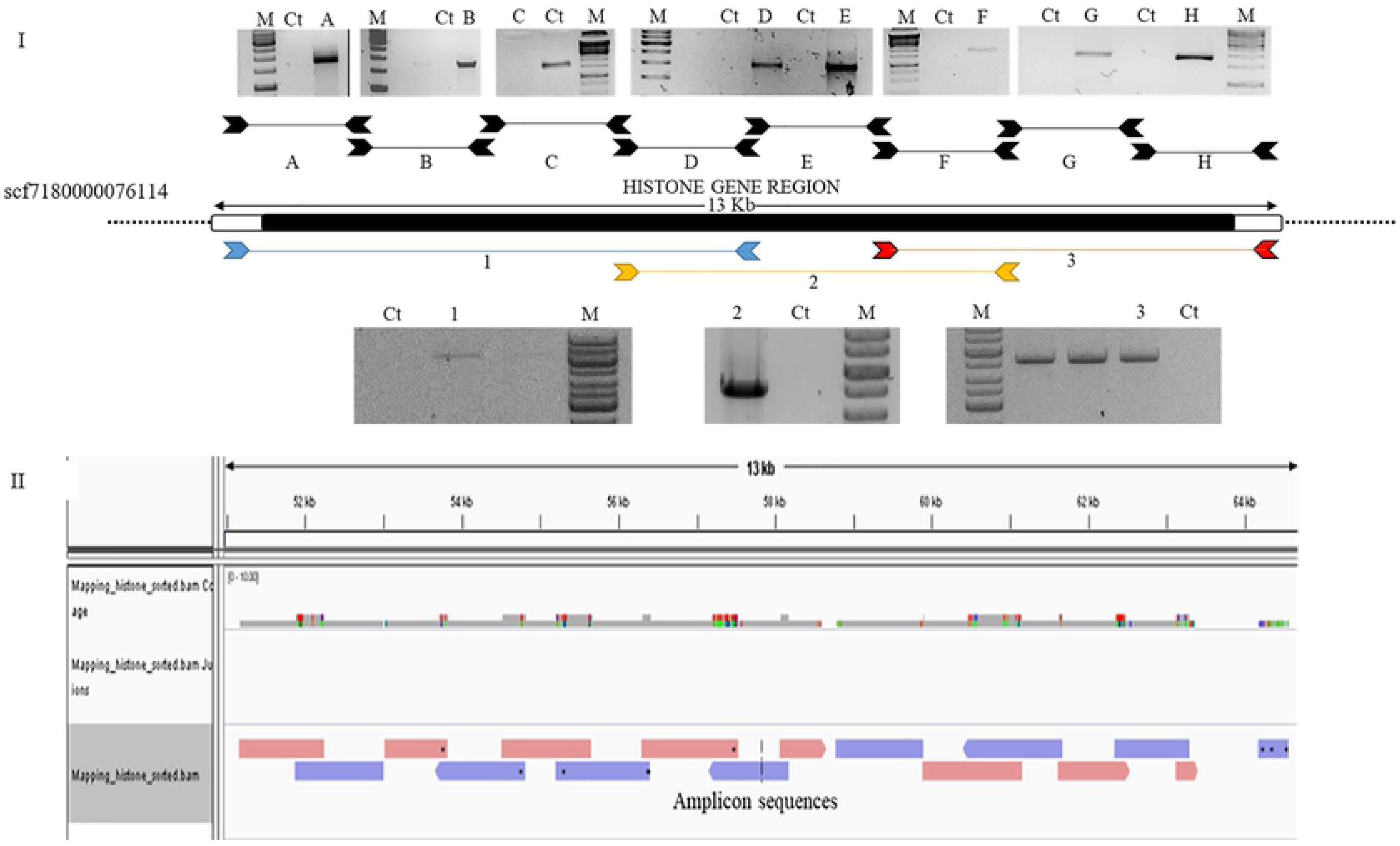
**Validation of Mhir genome assembly.** I. The amplicons obtained with the primer sets (A to H) used for PCR on scaffold, scf 0000076114. The double-headed arrows indicate the position of the forward and reverse primers while the coloured arrow-heads mark the position of the primers used in tiled long-PCR that map on the scaffold. The corresponding amplicons obtained are shown in gel images II. Alignment of sequences obtained by Sanger’s method of the long-PCR amplicons on Mhir genome assembly using Gene Viewer (IGV).

### Horizontal Gene Transfer (HGT) identification and Validation

Horizontal gene transfer (HGT) which refers to the lateral movement of genetic material between different species as opposed to direct descent, is very common in sap feeding insects. In addition, sap-feeders have obligate endosymbionts and this tripartite nested arrangement of obligate endosymbionts in mealybug, provide nutritional benefits to the host [17]. This may lead to lateral transfer of genes more frequently in those insects.

We analysed the HGTs in Mhir genome as well as reanalysed HGTs in *M. hirsutus* genome assembly provided by Husnik [18]. We identified 98 HGTs after applying all the QC criteria. These HGTs contain proteins involved in amino acid metabolism, vitamin biosynthesis and peptidoglycan metabolism (Fig3I). The identified HGTs were compared with the known HGTs reported by Husnik and McCutcheon [18]. In addition to those reported earlier, 29 novel HGTs coding for antioxidant enzymes, protease inhibitors, bacterial toxins and carbohydrate metabolism proteins were detected (Fig 3I). We validated five HGTs in Mhir using long PCR method with primers mapping in the adjacent host genes (arthropoda origin) and the HGTs (Fig 3II-IV).

**Fig 3.**
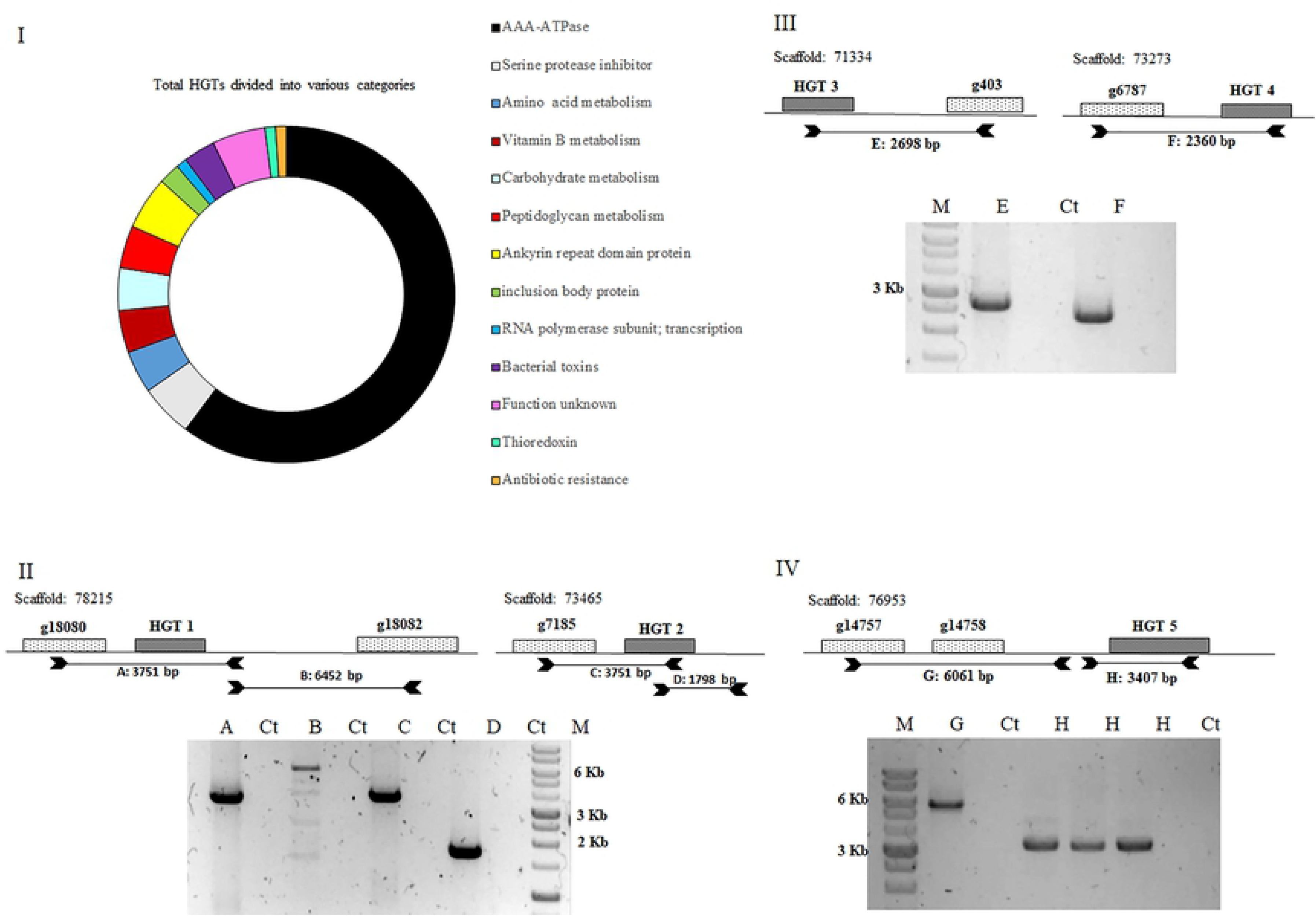
**Functional classification and validation of HGT in *Mhir* genome**. I-Donut plot of total HGTs classified into different functional classes; II, III & IV-Validation of HGTs by PCR amplification from Mhir genomic DNA. Genomic regions targeted as templates are indicated with the scaffold number; host genes (unfilled box) indicated with their gene Ids while HGTs (grey box) indicated as HGT 1, 2, 3. Primer position and the amplicons (A-H) are shown as double arrowed. M: 100bp marker; Ct: control without template DNA. HGT1: biotin synthase; HGT2: diamino pimelate epimerase; HGT3: dethiobiotin synthase; HGT4: AAA ATPase; HGT5:tryptophan 2-monooxygenase oxidoreductase. Host genes, g18080: nudix hydrolase 8; g18082: cytokine receptor isoform X2; g7185: peroxisomal acyl-coenzyme A oxidase 1; g403: cathepsin B; g6787: Uncharacterized protein; g14757: Retrotransposon protein; g14758: remained unannotated

Seven out of eight HGTs identified earlier are involved in amino acid and vitamin metabolism, while *tdcF* (reactive intermediate deaminase), involved in threonine metabolism, is detected only in our analysis. These HGTs are important as they participate in metabolic patchwork along with endosymbiont genes to complete the biosynthetic pathways of essential amino acids and vitamins [18]. Three bacterial genes involved in peptidoglycan metabolism were detected as HGTs, which may have a role in the maintenance of mutualistic relationship between the host and the endosymbionts. LD-carboxypeptidase (*ldcA)* and a rare lipoprotein A (*rlpA*) are HGTs in *A. pisum* and are absent in their endosymbiont, *Buchnera aphidicola*[19, 20]. Three bacterial toxin genes and an antibiotic resistant geneare among the HGTs in Mhir genome, which mayconfer resistance to bacterial pathogens. One such example is found in vinegar flies (Diptera: *Drosophilidae*) and aphids where, *cdtB* (cytolethal distending toxin B) coding for eukaryote-targeting DNAse I toxin is a HGT that confers resistance against parasitoid wasps [21]. Other HGTs identified in our analysis include AAA-ATPase, serine protease inhibitor, ankyrin repeat domain protein, inclusion body protein and thioredoxin. 91 out of the 98 HGT genes are expressed in both adult males and females while nine genes showed differential expression based on transcriptome analysis. All the nine DE HGTs exhibited higher expression in females (S2Table). The HGTs having differential expression included genes coding for protein degradation (AAA-ATPases), Vitamin B metabolism (*ribD, bioB*) and amino acid metabolism (*tdcF*). As all of them showed higher expression in females, suggesting their possible role in completing the metabolic network existing in mealybugs between host genes, β and γ-proteobacteria endosymbionts and HGTs for amino acids and vitamins biosynthetic pathways.

### Gene classes expanded and contracted in the mealybug genome

The evolutionary gain and loss of genes contribute to adaptation of organisms to their habitat. In insects like the mealybugs, the expansion and contraction of genes may be linked to their widespread geographical distribution and broad host range, thus it is interesting to analyse the gene classes that have expanded or contracted. To identify such gene classes, we compared the proteome of *M. hirsutus*and *P.citri* with five other insect species, namely*,D.melanogaster, A. pisum, R.prolixus, C.lectularius* and *B.mori* using OrthoFinder [22]. In this analysis, we identified orthogroups (cluster of orthologous genes) which were further classified as - Expanded, contracted and mealybug-specific based on the gene counts and a consolidated list of all the three clases is given in S3Table.

### Expanded gene classes; pesticide and desiccation resistance genes

Gene orthogroups involved in biological processes related to insecticide resistance, desiccation resistance, radiation resistance and hormone signalingare expanded in both Mhir and Pcit (Fig 4).

**Fig 4.**
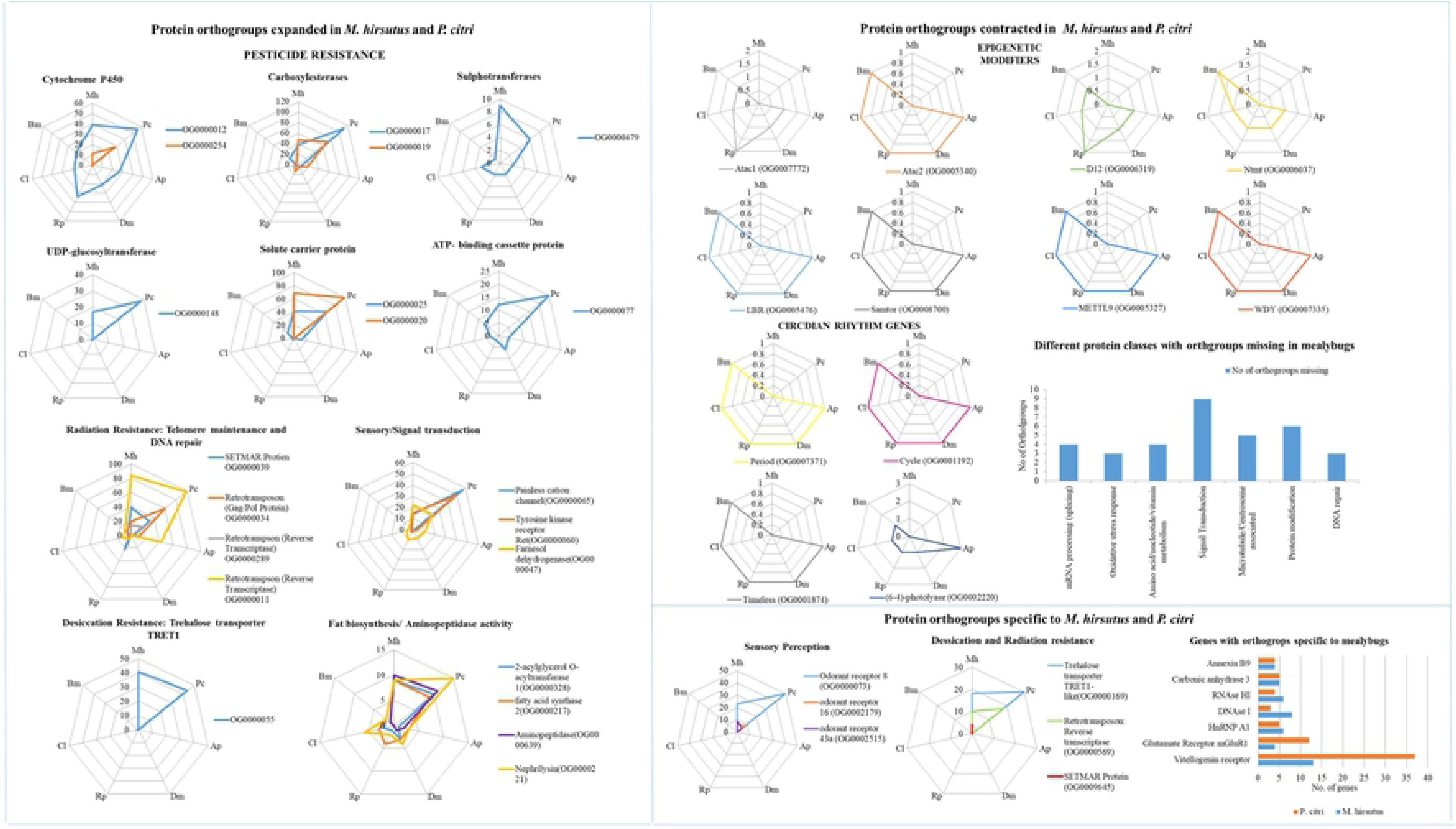
**Expanded and contracted gene classes in Mhir and Pcit**. The data derived from comparative analysis of protein classes using OrthoFinder, that are over-represented or under-represented in mealybug genome (Mh and Pc) relative to *A.pisum* (Ap), *C.lectularius* (Cl), *D.melanogaster*(Dm), *R.prolixus*(Rp), *B.mori*(Bm) is shown. The spider plots indicate the number of genes in the different species; A-Expanded, B-Contracted, C-Specifically found in mealybugs in the present comparative analysis. The functional class, such as pesticide resistance, radiation resistance is indicated at the top of each spider plot. The Orthogroup number and the associated function are indicated in each spider plot. The bar diagram under B shows the Orthogroups (not shown in spider plots) of different gene classes under-represented in mealybug genome and the bar diagram under C shows the Orthogroups found only in the mealybugs in our analysis (but not shown in spider plots). The complete list of genes specific to the mealybugs in our analysis is given in S3-S6 Tables.

The entire carboxylesterase gene family is expanded in the mealybugs, having the highest number of genes combining all the orthogroups of this gene family, Mhir has 104 and Pcit208 genes, while the number of genes in other insect genomes ranged from 20-40 (Fig4, S3 and S4 Table). Cytochrome P450 family also shows similar expansion, with mealybugs having the maximum number of genes in all the orthogroups put together, 95 genes in Mhir and 148 in Pcit, as compared to other insectsexcept *Rhodnius* (102 genes) (S3&S5 table). The Cytochrome P450 monooxygenases and Carboxylesterases enzymes are involved in first phase of insecticide detoxification, acting on a broad range of insecticides are expanded [23, 24],.One or more orthogroups of other genes associated with different phases of insecticide detoxification including phase II enzymes UDP-glycosyltransferases (UGTs) (Mhir-17, Pcit-38 genes) and Sulfotransferases (Mhir-9, Pcit-6 genes) and phase III enzymes, the ABC (ATP-binding cassette) transporters (Mhir-12, Pcit-25 genes) and solute carrier proteins are also expanded in mealybugs. *M. hirsutus* causesbrief but noticeable pest outbreaks, which can be attributed to its high reproductive potential, large brood size, high dispersal ability (at the crawler stages) and wide host range. The identification and expansion of the insecticide degradation and detoxification genes in the mealybug genome can be correlated with insecticide resistance known and hence their infestation in cultivated plants [25, 26]. In addition to the expansion of genes for insecticide metabolism, fatty acid metabolism enzymes, fatty acid synthase and acylglycerol-o-acyltransferase are also expanded in the mealybugs.These genes could be associated with the production of the waxy coating in mealybugs, majorly composed of trialkyl glycerols and wax esters. The waxy covering poses serious impediment for permeability of pesticides and protects mealybugs from desiccation and also predators [26, 27]. Thus, expansion of these genes could also contribute to resistance against insecticides.

The mealybugs, being universal pests,havethe ability to tolerate extreme environmental conditions. The trehalose transporter gene, Tret1-like, is another expanded gene class in the mealybug genome (Fig4 and S3Table). Trehalose is an important disaccharide that functions as a cryo-protectant, important for desiccation tolerance in insects and cannot enter the cells without a transporter[28].One Orthogroup of *Tret1* gene is expanded in mealybugs while another orthogroup of *Tret1* is represented as a specific class, present only in Mhir and Pcitand not in other genomes that we analysed. These findings suggest that mealybugs may show better tolerance to desiccation and may survive in xeric regionsas well.*SETMAR* and the retrotransposon proteins are the other major expanded gene classes, which are associated with DNA repair and telomere maintenance, respectively, that may contribute to radiation resistance in mealybugs [29, 30]. Further, one of the *SETMAR* orthogroups (OG0009645), containing 4 copies is specific to Mhir.

The other genes expanded in the mealybug are metallopeptidases (aminopeptidase, neprilysin) and farnesol dehydrogenase (S3Table). Neprilysins are M13 zinc metallopeptidase involved in reproduction in *Drosophila*and mammals [31]. Aminopeptidase Nis known to interact with insecticidal CryIA toxin (*Bacillus thuringiensis*) in lepidopterans and also participates in digestion and parasite vector interactions [32]. Farnesol dehydrogenase, another expanded class, is involved in biosynthesis of juvenile hormones that play essential role in reproduction, metamorphosis, development, polyphenism, and behavioral changes of insects and thus serve as good targets of insecticides [33, 34].

### Gene classes specific to Mhir and Pcit

Apart from expanded gene classes certain orthogroups are specific to mealybug genomes Mhir and Pcit, and are absent in other insect species chosen for comparison (S3Table). Gene orthogroups specific to mealybugs are mainly associated with olfactory sensation and oxidative stress/radiation resistance.

The orthogroups of proteins for chemosensory systems like odorant binding proteins, odorant receptors and olfactory receptors are not only expanded, but some are specific to mealybugs. These genes play an important role in the sophisticated olfactory system of insects through identification and binding of various odorants followed by signal transduction thereby affecting insect behavior [35]. The metabotropic glutamate receptor, specific to the mealybugs, is another gene associated with sensory perception. Considering the expansion of genes related to olfactory perception, one could speculate that these might help in detecting food sources and avoiding toxins. Some orthogroups of genes, carbonic anhydrase 3 and vitellogenin receptor are specifically present in the mealybug genome. Carbonic anhydrase 3 hasa protective role against oxidative stress [36] and may serve as an antioxidant, contributing to radiation resistance in mealybugs. Vitellogenin receptor is critical for oocyte development as it mediates suptake of vitellogenin [37].

### Contracted Gene Classes

The genes of circadian rhythm pathway are identified as contracted gene classes in Mhir and Pcit, with the genes period (*per*), cycle (*cyc*), timeless (*tim*), *CRY1 and CRY2* being absent. In contrast, the *Clk*, *Vri* and *Pdp1* genes are present (S1 Fig, S3Table). The circadian rhythm pathway in mealybug shares similarities with that of *Drosophila* and mammalian pathway (S1Fig). The absence of many of the core components of the circadian clock pathway, suggest lack of circadian rhythm in mealybugs, but several studies have suggested otherwise. Studies on flight activity in male mealybugs are dependent on the onset or exposure to light as well as an endogenous circadian rhythm [38]. The daily flight activity revealed that males of *P. citri*, *P. ficus* and *Ps.comstocki* are morning fliers while *M. hirsutus* and *N.viridis* (Newstead) fly around sunset time [38]. *Timeout* and *timeless* are two paralogous genes, considered to have originated by a duplication event, with mealybug containing only *timeout*. Hymenopterans like ants, bees and wasps similarly have the *timeout* gene, and have lost *timeless*. As in hymenopterans, timeout is under a strong positive selection in the absence of timeless and compensates for its function [39], similar functional substitution could also occur in mealybug. In the light of these observations, it is unclear whether circadian rhythm is operative in mealybugs.

Several epigenetic modifier genes are absent in the mealybug genome, these include components of ATAC histone acetyltransferase complex namely *Ada2a, Atac2, Atac1 and D12*. ATAC complex mediates acetylation of histone preferably H4 and is essential in *Drosophila melanogaster*. This complex containing two catalytic subunits i.e. *Gcn5* and *Atac2* plays important roles in signal transduction, cell cycle progression and facilitate nucleosome sliding catalysed by ISWI and SWI-SNF chromatin remodelers [40, 41]. Though ATAC complex components are absent in mealybugs (Mhir and Pcit), all other components of the complex (*Gcn5, Ada3, Hcf, wds, Chrac-14, NC2β, CG30390, Atac3*, and *Mocs2B*) are present. Considering the essential role of this complex both in Drosophila and mammalian development [40, 42], and the absence of its core components in mealybugs, it is possible that other HAT complexes might take over its role. The absence of these genes in both Pcit and Mhir largely rules out genome sequence error. Otherepigenetic modifiers missing in mealybugs include RNA methyltransferases *METTL9*, Samtor; histone methyltransferase *Ntmt* and WD40 repeat protein *WDY* (part of Set1/COMPASS complex). Apart from epigenetic modifiers, some of the genes for DNA repair, oxidative stress response belong to the contracted class and the mode of compensation for such functions needs to be investigated (S6Table).

### Analysis of selected gene classes in mealybug genome

#### HOX gene clusters in the mealybug genome

The homeotic (Hox) genes form a distinct class oftranscription factors, belonging to the homeobox gene superfamily, involved in the cell fate decisions during development and are highly conserved[43]. In the Mhir genome, except *Antp* and *Scr* which are single copy genes, all other hox genes are present in 2 or more copies compared to the other insect species (Fig 5). The hox genes in Mhir lack the cluster-like arrangement seen in *Drosophila* or *Tribolium,* and are scattered throughout the genome with some genes present in pairs or triplets on the same scaffold, while some are on the same scaffold but interspersed with other gene classes (Fig 5). This kind of arrangement of hox genes is called the “atomized or no clustering” which is also observed in flatworm *Schistosoma mansoni* and nematodes[44]. *Anopheles gambiae*, *Tribolium castaneum* and *Cimex lectularius* have a single large cluster of all hox genes, *Drosophila* and *Bombyx mori* consists of split cluster with hox genes divided between the two sub-clusters [44, 45].

**Fig 5.**
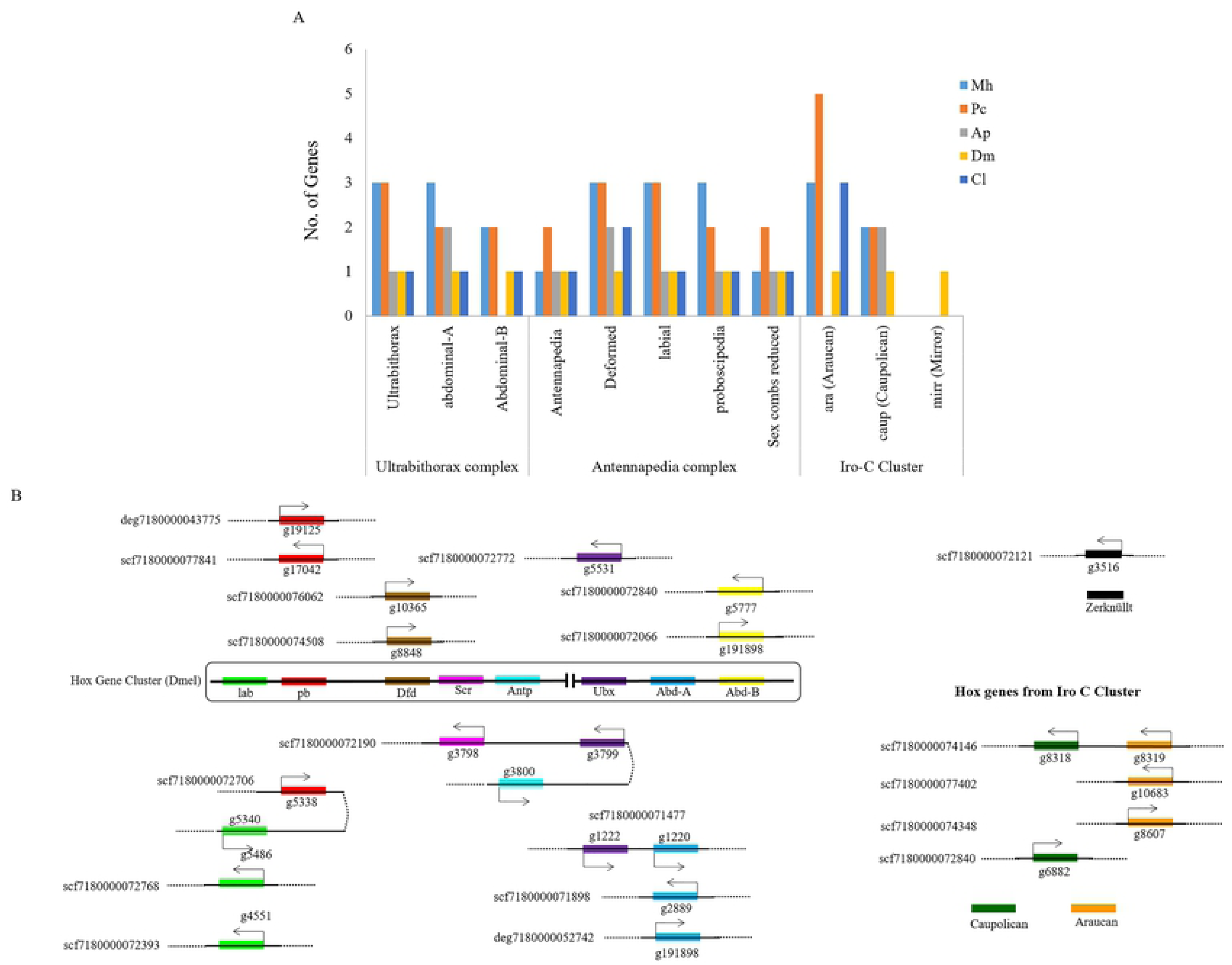
**Homeotic (Hox) genes in *Mhir* and *Pcit* genome**. A-Copy number of genes of different Hox clusters compared with that of other insect genomes. B-Line diagram representing the relative position of the Hox genes in different scaffolds of *Mhir* genome. Ultrabithorax, Antennapedia and Iro-C complex of Drosophila (Dme) used as reference are shown within the boxes. Mh: *M. hirsutus*; Pc: *P. citri*; Ap: *A. pisum*; Cl: *C. lectularius*; Dm: *D. melanogaster*; Rp: *R. prolixus*; Bm: *B. mori*. The figure is not drawn to scale.

Iroquois-family of genes are another conserved group of homeodomain containing transcription factors which play a major role in defining the identity of large and diverse territories of the body such as the dorsal region of head, eye and mesothorax in *Drosophila*. They are usually present as one or two clusters of three genes. *Drosophila* hasthree genes belonging to Iroquois family-*mirror (mirr), araucan (ara)* and *caupolican (caup)* which together form Iroquois-Complex (Iro-C) [46]. In Mhirgenome only two members of the Iro-C family, *ara* and *caup* are present while the *mirror* gene is absent. Benoit et al,[45] reported the presence of two Iro-C genes in *C. lectularius, mirror* and *Iroquuois (Iro)* which they found to be orthologous to tandem paralogs of *araucan* and *caupolican* of *Drosophila*, though we failed to find *mirror* gene in mealybugs.

#### Identification of epigenetic modifiers in mealybug genome; retrieval and analysis of functional classes

In the absence of sex chromosomes, sex determination in mealybugs is very closely correlated with genomic imprinting. We have mined the components of the epigenetic machinery in Mhirand Pcit, as important players in developmental regulation and also the differential regulation of homologous chromosomes. The machinery for methylation of DNA and histones is analysed considering the presence of the writers, readers and the erasers of epigenetic marking of the genome.

The genes coding for histones, the primary substrates of epigenetic modification, and their variants were identified in the mealybug genome along with other genomes for comparison (Fig 6A, B). Mhir contains only a single complete quintet cluster of histone genes, while in two of its other quintets, the histone H1 is absent (Fig 6C). The remaining histone genes are present in scaffold either singly or with some of the other histone genes, but not all the histones. For instance, H2A may be present as an isolated gene in one scaffold or in combination with some of the core histone genes in another scaffold (Fig 6C). This organization has some similarity with the other hemipteran, *Acyrthosiphon pisum* [47]. The number of alleles for core histones vary with only two copies of histone H1 gene present in Mhir while 9 copies in Pcit.

**Fig 6.**
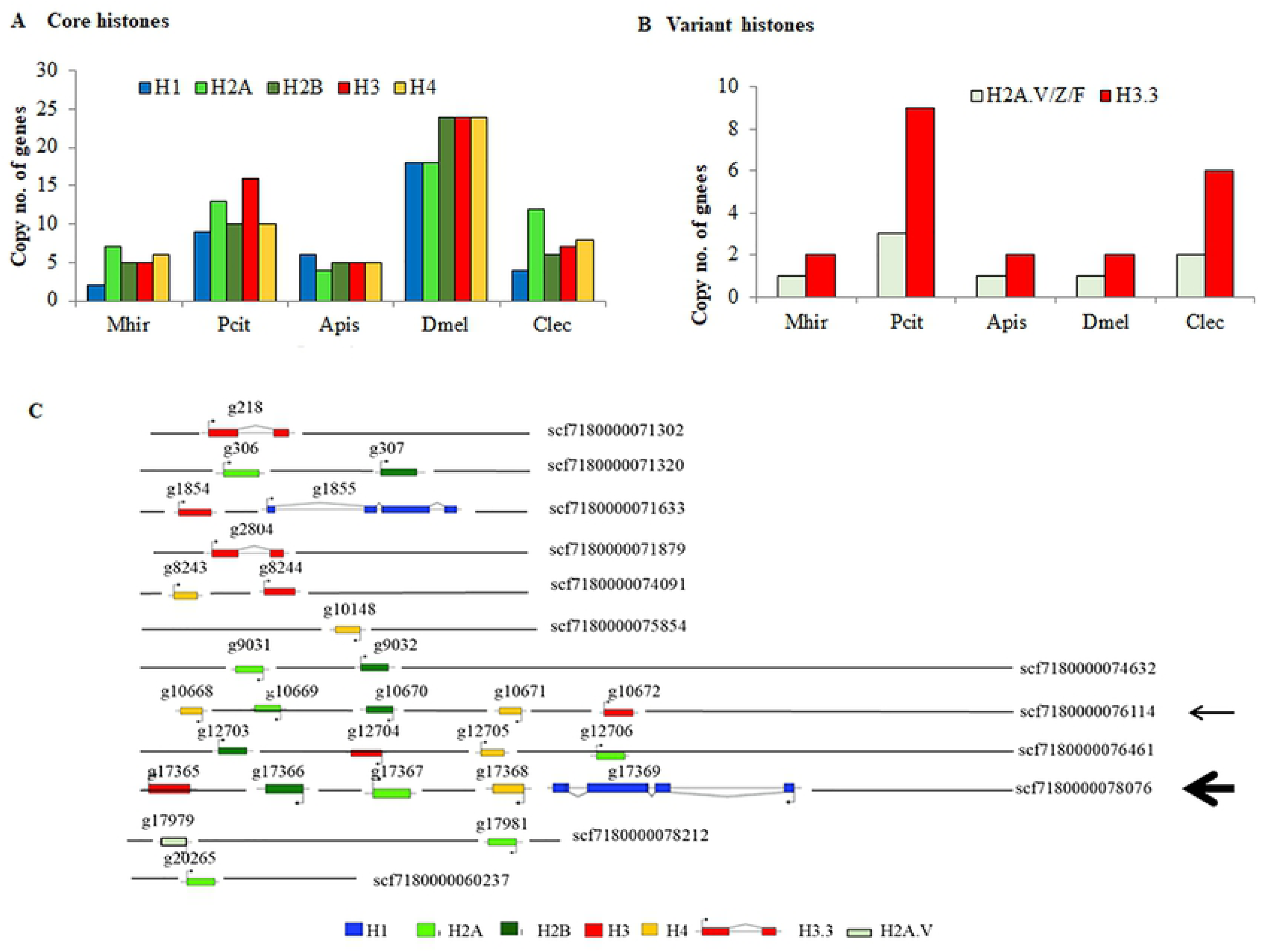
**Histone and variant histone genes in Mhir**. The copy number of core histone genes (A) and variant histones (B) in Mhir genome is compared with that of other insect genomes. C-mapping of histone clusters on Mhir genome. The numbers written with prefix scf or deg are the scaffold/contig IDs. scf7180000078076 (marked with a thick arrow) contains the complete quintet cluster while scf180000076114 and scf180000076461 have histone H1 gene missing from the quintet clusters. scf7180000076114 has been used for validation of assembly (thin arrow). The figure is not drawn to scale.

MhirandPcithave multiple copies of the genes for the variant histones H2A.V and H3.3. The histone H2A.V is required for heterochromatin assembly and DNA damage response in *Drosophila*[48]. Mhir has two copies while Pcit has nine copies of histone H3.3, which is evolutionarily conserved and is associated with pericentromeric and telomeric regions where it replaces the canonical histone H3 during transcription [49].

#### DNA methylation machinery in mealybug

DNA methylation is associated with several epigenetic phenomena like genomic imprinting, X-chromosome inactivation and transposon repression in mammals [50]. It is one of the key molecular mechanisms associated with mammalian imprinted genes containing differential methylated regions (DMRs) which are methylated in parental-origin specific manner [51, 52]. Differential methylation of the genome in males and females is detected in different species of mealybugs [11, 14]. The major DNA modification associated with imprinting and other epigenetic processes is cytosine methylation (5mC), predominantly at CpG dinucleotides; however, non-CpG DNA methylation has also been reported [14, 50]. We analyzed the DNA methyltransferases and demethylases in Mhir and Pcit genome and compared them with those in other genomes (S7Table). Since,*Drosophila* lacks canonical DNA methyltransferases 1 and 3 homologs [53], we included human DNMT proteins as reference for comparative analysis. We found two types of DNA methyltransferases: cytosine-specific and adenine-specific DNA methyltransferases in all the insect species analysed. These methyltransferases contain S-adenosyl-L-methionine-dependent methyltransferase (IPR029063) as the functional domain while adenine specific DNA methyltransferases, contain an additional signature IPR007757 representing MTA-70-like protein family. Multiple copies of cytosine methyltransferase *DNMT1* genes are found in *P. citri,* while *M. hirsutus* has a single copy.. The adenine DNA methyltransferase, *METTL4* gene is present as a single copy in both. We also generated phylogenetic trees of DNMTs of all insects to assess their evolutionary conservation with human proteins (Fig 7).

**Fig 7.**
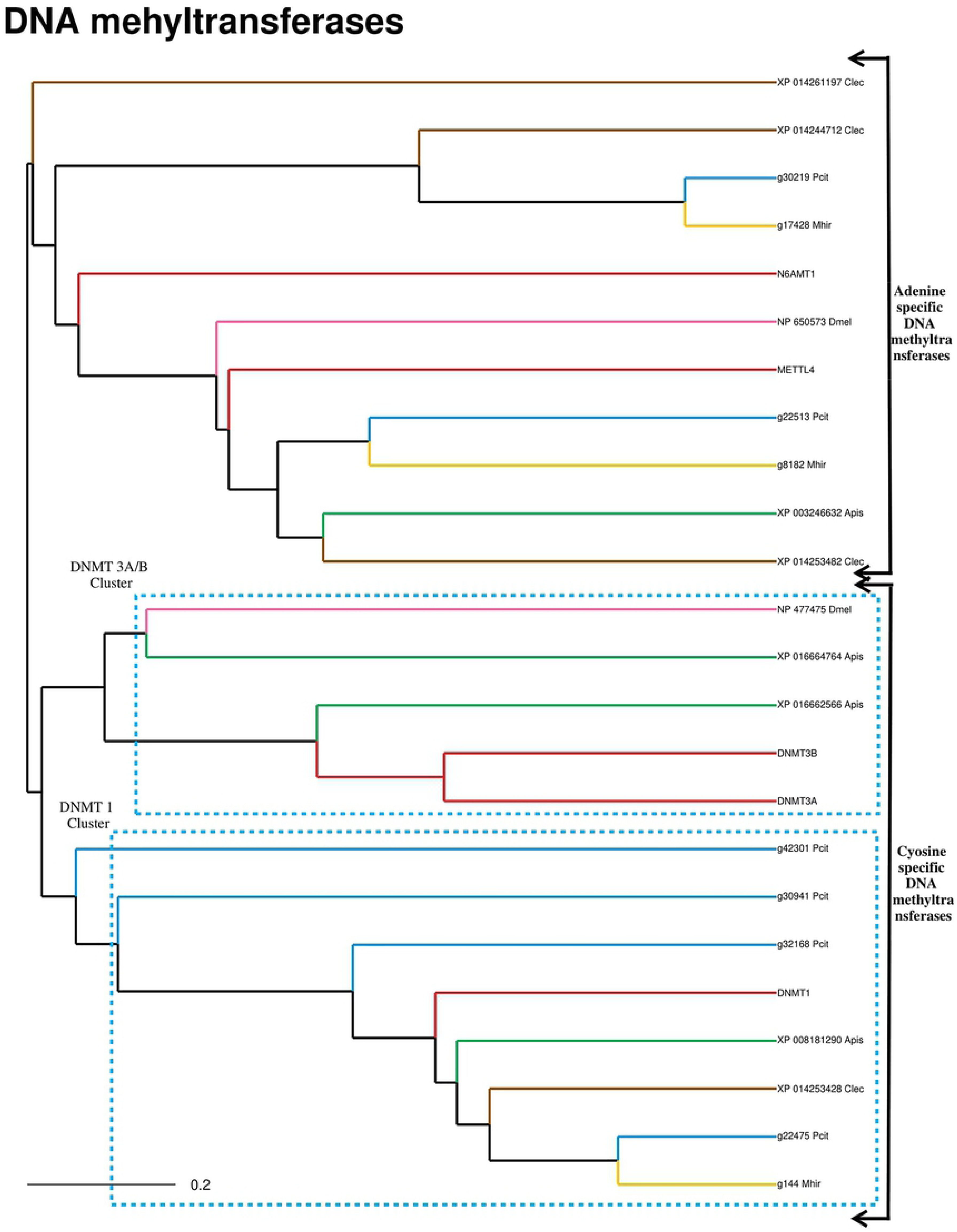
**Phylogenetic clustering of DNA methyltransferases (DNMTs) based on multiple sequence alignment of Mhirwith other genomes**. Human DNMTs were used as reference. Adenine specific DNA methyltransferases (N6AMT, METTL4) and cytosine specific DNA methyltransferases (DNMT1 and DNMT3) cluster separately. Cytosine DNMTs cluster further divides into two subclusters of De novo methyltransferases (DNMT3A, 3B) and maintenance methyltransferase (DNMT1) (shown by dotted line). The key for the colour code is given as inset.

We observed that DNMT proteins of *M. hirsutus, P. citri* and *C. lectularius* and *A. pisum* clustered with DNMT1, while only one methyltransferase of *A. pisum* clusters withDNMT3. *P. citri* proteins (g3941, g42301), remained outliers, as they are partial sequences lacking functional domains. Except for *A. pisum,*the *de novo* methyltransferases are absent in all the other insects including the mealybugs.

Though mealybug lacks DNMT3, the presence of cytosine DNA methylation was established in *Planococcus lilacinus*[14]. The study estimated the frequency of occurrence of 5mC in CpG as well as other dinucleotides and found that 5mC in CpG, CpA and CpT dinucleotides occurs at comparable frequency while the frequency of CpC methylation is lower [14]. Bewick et al, [54] showed the loss of *DNMT3* and presence of only maintenance methyltransferase genes in several insect species which nevertheless have DNA methylation. These findings suggest that DNMT3 may be expendable for DNA methylation or DNMT1 could compensate forDNMT3. In the light of these observations, we compared the domain architecture of DNMTs in Mhir andPcit with that of humans. DNMTs ofMhir and Pcitcontain all the domains associated with DNMT1 along with C-5 cytosine-specific DNA methylase domain essential for function, but lacked the characteristic PWWP domain of *DNMT3*.

We found DNA methyltransferase for 6mA in Mhir, Pcit and other insects which clustered with human METTL4. Another group of proteins formed a separate cluster with human N6AMT1 protein, a 6-methyladenine-specific DNA methyltransferase later shown to be involved in methylation of elongation factor protein, suggesting dual substrate specificity[55]. Greer et al, [56] [43]detected 6mA DNA modification in *C. elegans* and demonstrated its role in trans-generational epigenetic inheritance. They identified 6mA methyltransferase, Damt-1 along with demethylase NMAD-1 in *C. elegans.* Their work suggested a cross-talk between DNA and histone methylation [56].

The presence of 6-methyl adenine (6mA) along with the genes for methyltransferase and demethylase (DMAD) is reported for Drosophila [57]. Demethylation of 6mA in transposon leading to repression demonstrated in Drosophila, implies that 6mA is a marker for active state [57]. The presence of methyltransferase (N6AMT1) and demethylase(ALKBH1) for adenine methylation in humans and the enrichment of 6mA in exonic regions in the genes is known [58]. Thus the identification of genes for methylation and demethylation of adenine in both Mhir and Pcit suggests a novel mechanism of differential enrichment of 6mA in females contributing to imprinting in mealybugs. The low density of 6mA in human X and Y chromosomes which have high 5mC levels is noteworthy [58]. The presence 6mA and 7mG has been demonstrated in mealybugs by Chandra and co-workers in *Planococcus lilacinus*[10]. DNA methyltransferase that could methylate poly (dC-dG) as well as poly (dC-dA) was also reported in *P. lilacinus*[59].

#### DNA demethylation machinery in mealybugs

We analysed Mhir, Pcit and other insect genomes for 5-methylcytosine- and 6-methyladenine-specific demethylases. In humans, TET and ALKBH are two major protein families that participate in demethylation of 5mC and 6mA DNA modifications, respectively [58, 60, 61]. Using BLASTp analysis, we identified genes for both the protein families in all insects. TET proteins are methylcytosine dioxygenases which regulate global levels of 5-methylcytosine and/or 5-hydroxymethylcytosine through active DNA demethylation [53, 60]. The TET protein family has 3 members: TET1, 2 and 3, all containing catalytic C terminal domain called 2OGFeDO, oxygenase domain (IPR024779), with TET1 and 3 containing additional N terminal Zinc finger, CXXC-type domain (IPR002857).

In humans, ALKBH1 is identified as 6mA demethylase [58]. The AlkB family of proteins are Fe(II)- and α-ketoglutarate-dependent dioxygenases that perform alkylated DNA damage repair through oxidative dealkylation [62]. Apart from DNA repair, these enzymes are also implicated in nucleotide demethylation, with ALKBH1 and ALKBH4 majorly associated with removal of 6mA DNA modification [58, 61, 63]. We generated a phylogenetic tree of DNA demethylases including human DNA demethylases and identified 3 major clusters (Fig8); two clusters containing human ALKBH proteins and a third big cluster C, for human TET proteins. Within this cluster, the human TET proteins formed a separate branch and the others segregated into two groups, one with Mhir, Pcit, Dmel TET proteins clustering together (Group I) and the other with N6 adenine DNA demethylases proteins from all insects (Group II). It was interesting to note that proteins identified as N6 DNA demethylases in BLASTp clustered with the TET protein family. To further dissect this issue, we compared the domain architecture of proteins forming two different branches but observed no significant difference as all proteins contained 2OGFeDO, oxygenase domain (IPR024779) and Zinc finger, CXXC-type domain (IPR002857) with the exception of 3 proteins g79 (Pcit), g6287 (Mhir) and XP_008183448 (Apis) which are partial sequences containing only Zinc finger, CXXC-type domain (IPR002857). Multiple sequence alignment with human TET proteins revealed that Group I proteins shared higher sequence similarity with human TET proteins than Group II proteins as shown in percentage identity matrix (S8Table), suggesting that group II proteins shared similarity only in domain regions and are variable in regions outside the domains. As mentioned earlier the group II proteins were identified as N6 DNA demethylases, could indeed be orthologues of TET proteins that serve as 6mA demethylases, as demethylation of 6mA in *Drosophila* is regulated by its TET homolog [57].

**Fig 8.**
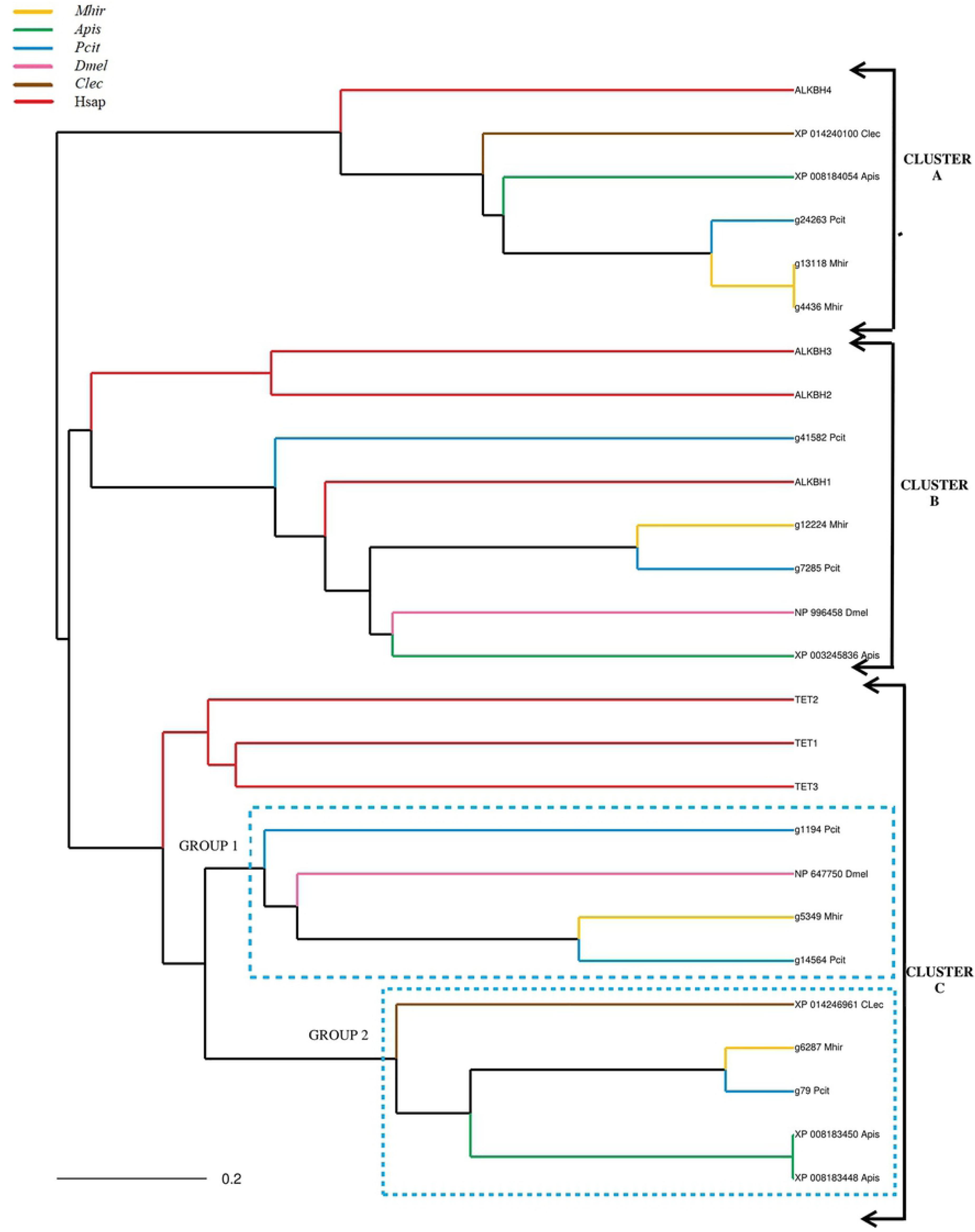
**Phylogenetic clustering of DNA demethylases by multiple sequence alignment**. Human DNA demethylases were used as reference. The three clusters formed: A-ALKBH4 Adenine demethylases, B-Adenine demethylases including human ALKBH3, ALKBH2 and ALKBH1 along with other insect ALKBH1 proteins, C-TET proteins (Cytosine specific DNA demethylases). The two sub-clusters under C segregate human TET proteins, while other having Mhir and Pcit is divided into two groups 1 and 2 (shown by dotted line).

The ALKBH family formed two clusters in the phylogenetic tree with one cluster (cluster A) containing ALKBH4 proteins of all insects grouping with human ALKBH4 protein and other cluster (cluster B) containing alkbh1 proteins of all insects grouping with human ALKBH1. Both ALKBH1 and ALKBH4 proteins shared the Alpha-ketoglutarate-dependent dioxygenase domain (IPR027450) while ALKBH4 proteins contained an additional Oxoglutarate/iron-dependent dioxygenase domain (IPR005123). ALKBH2 and ALKBH3formed a separate cluster; we could not find orthologs for these proteins in Mhir and Pcit as well as in other insect species.

#### Mining the genes for epigenetic modification of histones

The various classes of genes were curated based on the presence of functional signatures or domains using InterProScan (https://www.ebi.ac.uk/interpro) from the annotated genome of *Drosophila melanogaster* as reference for the newly sequenced genome of Mhir. Based on the frequency of occurrence of the domains in different genes of a given functional class in the genome of *D.melanogaster*, we selected domains which we designate as high priority domains (HPD; [64]). In a similar analysis, the SET and Pre-SET domains were identified as high priority domains for histone methyltransferase as described earlier [64].

In-house PERL script was used to fetch genes containing high priority domains in Mhir and also in the other genomes that were used for comparative analysis. The gene classes were also mined using BLASTp (www.blast.ncbi.nlm.nih.gov).After manual curation, these genes were divided into three groups-(i) Interproscan only - genes which harbor the HPD but has not been annotated/assigned any function in BLASTp (ii) BLASTp only-those genes that lack HPD even though a function is assigned in BLASTp (iii) Concordant-those that are annotated by BLASTp and contain the HPD as well (common to both InterProScan and BLASTp). A representative profile for histone acetyl transferases (HATs) shows the Acyl CoA acyl transferase and histone acetyltransferase_MYST_type domain as the high priority domains. The Chromatin remodelling proteins have four high priority domains viz. Helicase ATP-binding, SNF2_N, Helicase-C and the P-loop NTPase (S2Fig). A similar criterion was utilized to identify the different classes of histone modifiers.

In all the genomes, there are a few genes having the high-priority domain(s), but not annotated in BLASTp. We consider these as potentially novel genes. The number of genes in this category is least in *C. lectularius* (8), while it is high in MhirandPcit at 26 and 12.In Mhir (71) genes are predicted to be histone modifiers by BLASTp only, lacking the high-priority domains (Fig 9A; S9Table). On manual curation, we find that these are partial sequences.Pcit has a significantly larger genome compared to Mhir andhas a higher number of almost all the genes that we analysed.

**Fig 9.**
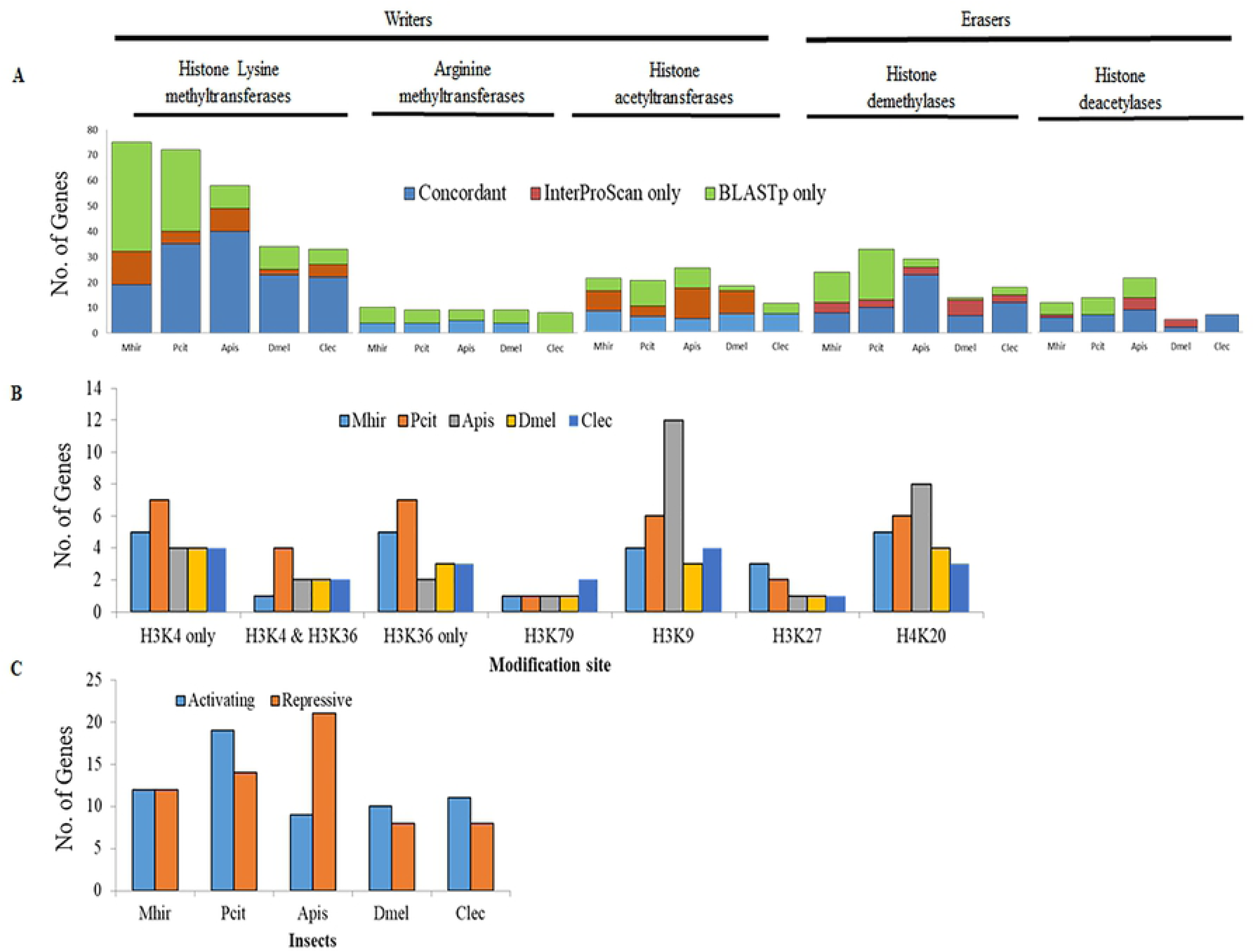
**Comparative analysis of the histone modifiers of Mhir and Pcit genome with other insect species**. A-The numbers of genes for the writers and erasers are compared. The genes identified by BLASTp only did not have the high priority domains (HPD), those identified by InterProScan only had HPD, but were marked as hypothetical/unknown in BLASTp, Concordant classes were annotated by BLASTp and InterProScan. The genes identified by InterProScan only are the potential novel genes. B-Total number of lysine histone methyltransferase genes for a specific modification in Mhir compared with other genomes, C- comparison of the number of genes under activating and repressive classes of histone lysine methyltransferases.

### Writers of epigenetic imprint

#### Histone methyltransferases (HMT)

The modification of histones is one of the major and better analyzed epigenetic markings [65]. The SET and Pre-SET are the high-priority domains for HMTs [64]. All the three categories, SET proteins, DOT1 proteins and the arginine methyltransferases involved in histone methylation were annotated in the Mhir and Pcit genome. Mhir has 19, Pcit 35 while *A. pisum* with 40 genes has the maximum number of histone lysine methyltransferase genes (Fig 9A, S9Table). Besides this, many genes with HPD are recognized (Mhir-13, Pcit-5, Apis-9), but were not annotated in BLASTp(Fig 9A). These are potentially novel methyltransferases, to be investigated further.The number of genes for specific modification as well as activating and repressive modifications is not high in mealybugs relative to that in others though a large part of genome is under epigenetic regulation(Fig 9B and 9C). This is not unexpected considering that these are catalytic functions.

We used phylogenetic clustering with *Drosophila* HMTs, to assign specificity of the coded enzymes (Fig 10). The low confidence annotations (identified by BLASTp only) are due to partial sequences, but the identity of some of them could be deciphered based on their clustering pattern. For example between the two E(z)-like genes, Mhir_g5597 is complete and Mhir_g18633 is fragmented, similarly additional copies of genes for trr detected (Mhir_g13137 & Mhir_g20142) are also partial sequences (Fig 10, S3A, B, C, D Fig). The key-word based identification of BLASTp annotated genes also led to error as in the case of trl and Ash2, which was detected by phylogenetic clustering (Fig 10). These proteins are not catalytic histone modifiers in *Drosophila*, but are associated with epigenetic regulatory complexes, like COMPASS containing histone methyltransferase for H3K4 trimethylation, while trl codes GAGA protein [66, 67]. There are multiple genes coding for almost all the methyltransferases in both Mhir and Pcit (Fig 10, S10Table).

**Fig 10.**
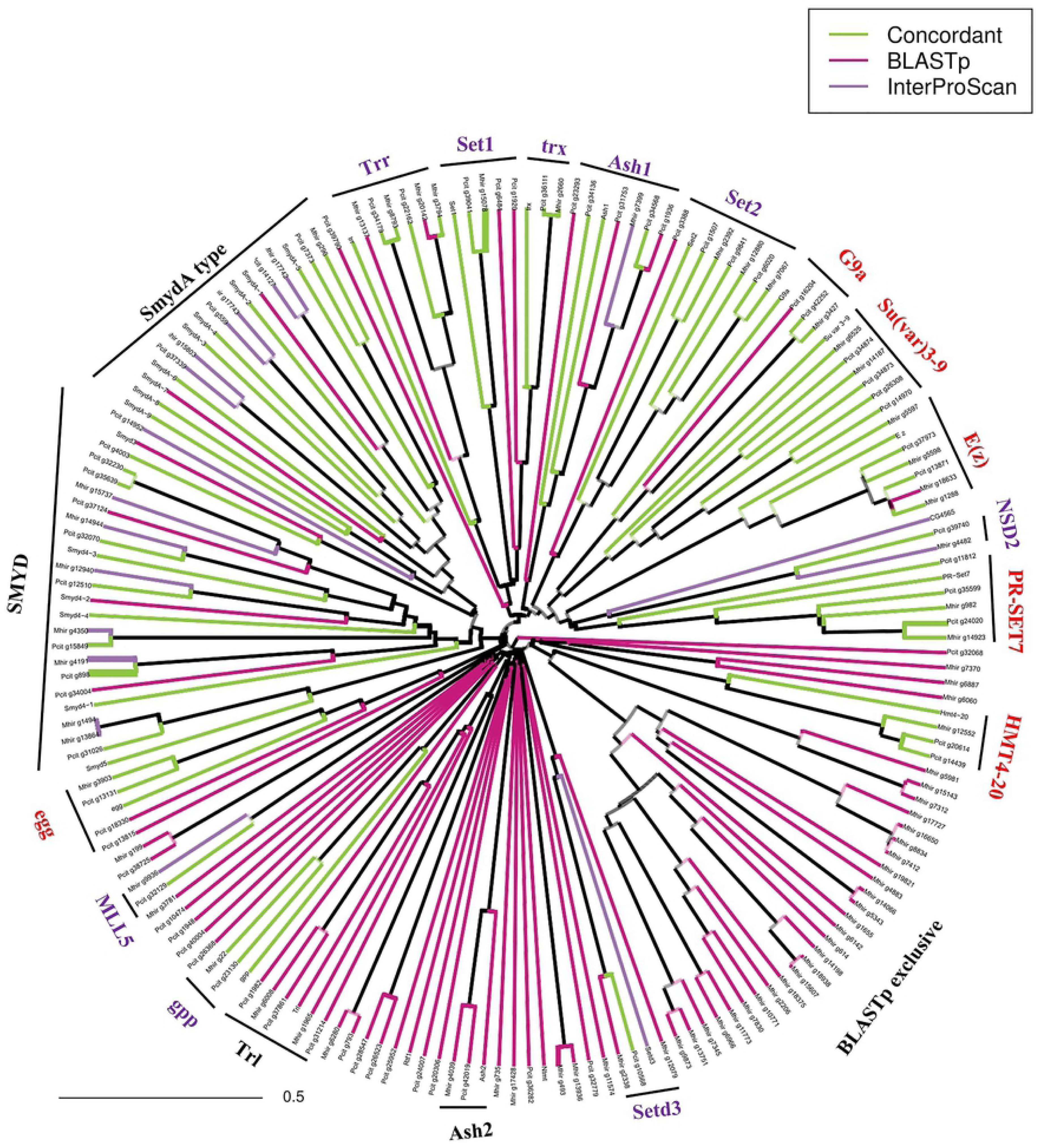
**Phylogenetic clustering of the histone methyltransferases (HMTs).** The protein sequences of HMTs of *Mhir and Pcit,* were aligned with those of *Dmel* as the reference sequence. The three classes (BLASTp only, InterProScan only and Concordant), are indicated by differently coloured lines, as given in the inset. The black lines outside the tree, indicate the proteins clustering with a known Dmel proteins, confirming Mhir and Pcit functional identity. The proteins from Mhir and Pcit have the suffixes Mhir and Pcit respectively, while the Dmel proteins are named according to the nomenclature in Uniprot. The activating (purple) and repressing (red) HMTs are indicated. Most of the putative novel methyltransferases cluster with SMYD proteins of Drosophila.

The methyltransferases are involved in the regulation of several pathways in almost all eukaryotes including yeast (S10Table). Their function is mediated through different multiprotein complexes that determine the site of action and the target genes. Among the various pathways modulated by histone methylation, those that result in silencing of genes and whole chromosomes are the most relevant in mealybugs.

The silencing histone methylation marks (H3K9me2,me3) are enriched in heterochromatin in nuclei from mealybug males and retained during spermatogenesis and further in the male pronucleus in Pcit [68]. Therefore it was considered as a candidate for male specific imprinting mark [68]. In Drosophila, there are multiple methyltransferases mediating H3K9 methylation: egg(SETDB1), G9a and Su(var)3-9 [69]. All the three genes were annotated in mealybugs (S10Table). The function of egg(SETDB1) is important for maintenance of heterochromatin in Drosophila and a balance between egg and Su(var)3-9 genes is essential asmutation in both the two genes is less deleterious than single gene mutation ([70]). This suggests an interaction between the two genes and in this context the higher copy number of genes in mealybugs is of significance. Their tissue specific and developmental stage specific expression has to be explored.

Similarly, there are two genes Pr-SET7 and the Suv4-20 that catalyse H4K20 modification, yet another repressive histone mark (Fig10). There is a functional cooperation between these two enzymes; Pr-SET7 catalyses the H4K20 monomethylation and Suv4-20 brings about trimethylation of H4K20 in *Drosophila* [71, 72].H4K20 methylation is important for chromatin condensation (during interphase) and position-effect variegation in Drosophila [73]. In mealybugs, H4K20 methylation is localized on the heterochromatized paternal chromosomes in males and in females no specific distribution was observed [12]. Mathur et al,[13] reported the presence of H4K20 in both male and female mealybugs, it was enriched in soluble chromatin fraction, that was not associated with nuclear matrix. Heterochromatin protein 1 (HP1) is essential for maintaining the normal levels of H4K20me3 [72, 74]. The identification of multiple copies of several of these genes in the mealybugs suggest shared components between constitutive heterochromatin seen in all organisms and the facultative heterochromatization observed in mealybugs and the inactive X chromosome of mammals. H3K36 methylation antagonizes Polycomb silencing and has a role in DNA repair and mRNA splicing [75]. There are multiple enzymes that bring about H3K36 methylation, among which Set2 is the only enzyme responsible for trimethylation of H3K36 [76]. Ash1 is H3K36me2-specific methyltransferase associated with trithorax complex and is also required for H3K4 methylation [76–78].The Ash1 and Set2 genes are well conserved in mealybugs (Fig 10, S4C Fig). MLL5 also methylates H3K36 in mammals, Mhir and Pcit is similar to mammalian MLL5([79]; S10 Table). SETMAR genes, identified as an expanded class of genes in mealybugs dimethylate H3K36 and are important for DNA repair activity [29].

SMYDs, bring about lysine methylation of non-histone proteins and some of them methylate histones also [80]. Among them SmydA genes are specific to arthropods [81]. There is evidence for the role of Smyd5 controlling expression of proinflammatory genes through histone methylation with the interface of the NCoR complex in mammals [82]. Both Mhir and Pcit have Smyd genes and Smyd5 is identified as a differentially expressed gene (Fig 10). A single ortholog of gpp/DOT-1 (H3K79 specific) is present both in Mhir and Pcit (Fig 10). DOT1L associated with the Dot complex is important for DNA damage response [83–85]. H3K4 methylation is brought about by the Set1, Smyd1,trr and the trx genes [86–90]. The Mhir trithorax-related (trr) has PHD finger domains and a HMG box (high mobility box domain) similar to the human MLL3 ortholog and not of Drosophila, where lpt and trr are two separate genes [91]. Mhir has one copy as a composite gene with both lpt and trr in addition to another where the two are coded by separate genes. The alignment of the trr gene in various insects is shown in S3C, DFig. The trr genes are highly conserved (Fig10; S3C, D Fig). In Apis, Pcit and Clec, the composite gene resembles the human homologue (MLL3). The bootstrapped trees for the lysine methyltransferases are given in S4 Fig.

Arginine methyltransferases are also present in Mhir (10 genes) and Pcit (9 genes, S9 Table). Most of the genes of a given function cluster together but a few are clustering away from their paralogues. Art9 of Mhir and Pcit are clustering with Art7 and they are far from the Drosophila Art9 showing sequence variability between dipteran and hemipteran Art9 (S5Fig). The detection of various modifications of histones reported in mealybugs, the detection of multiple genes coding for the enzymes facilitates their tissue and stage specific expression studies. In the absence of sex chromosomes in mealybugs, epigenetic mechanisms are important part of sex determination and differential regulation of the homologous chromosomes.

#### Histone acetyltransferase

The acetylation of histones is widely encountered in almost all eukaryotes and is associated with transcriptional activation and open chromatin state. The HATs (Histone acetyltransferases) are divided into two superfamilies, the MYST-type and Gcn5-related. As described under the methods section, the genes were annotated using BLASTp and InterProScan in Mhir and Pcit along with other species. The phylogenetic analysis of these proteinsled to the formation of a super-cluster comprising of MYST-type HATs namely MOF, Tip60, CHM, ENOK and CG1894 along with a number of Mhir and Pcitproteins (Fig11). The MOF sub-cluster includes two Mhir proteins (g14997 and g16369) along with Pcit genes (g7142, g40821). MOF is an H4K16-specific acetyltransferase that participates in dosage compensation in *Drosophila* [92, 93]. The H4Ac16 leads to hyper-transcription of X-chromosome in male flies whereas the female X-chromosome shows decreased enrichment of H4Ac16 [94]. The differential enrichment of H4 acetylation on active maternal and the heterochromatic male genome in Pcitis reported earlier ([95]).

The Mhir proteins (g9382 and g9310) and Pcit protein (g36521) cluster with Dmel Tip60, which is closely related to MOF and participates in acetylation of histone H4 and H2A ([96, 97]; Fig.11). Mhir g4136 and Pcitg39105 cluster with Dmel CHM and ENOK proteins, which acetylate histone H3 and are involved in position effect variegation. The haploinsufficiency leading to a phenotype indicates that the dosage of this gene is also critical [98]. Thus, the physiological haploidy resulting from the inactivation of the paternal genome in mealybugs might be correlated to differential histone acetylation. Chm which is involved in PcG mediated silencing, clustered in a super-cluster that included MOF, Tip60, CHM, ENOK and CG1894, ([98, 99] [93, 94],Fig11). Mhir proteins g14997 & g16369 along with Pcit proteins g7142 & g40821 that cluster with MOF harbour Chromo-like domain. Though, the length of the homologous proteins in Mhir and Pcit is reduced, they are similar to the C-terminal regions of Dmel protein.

**Fig 11.**
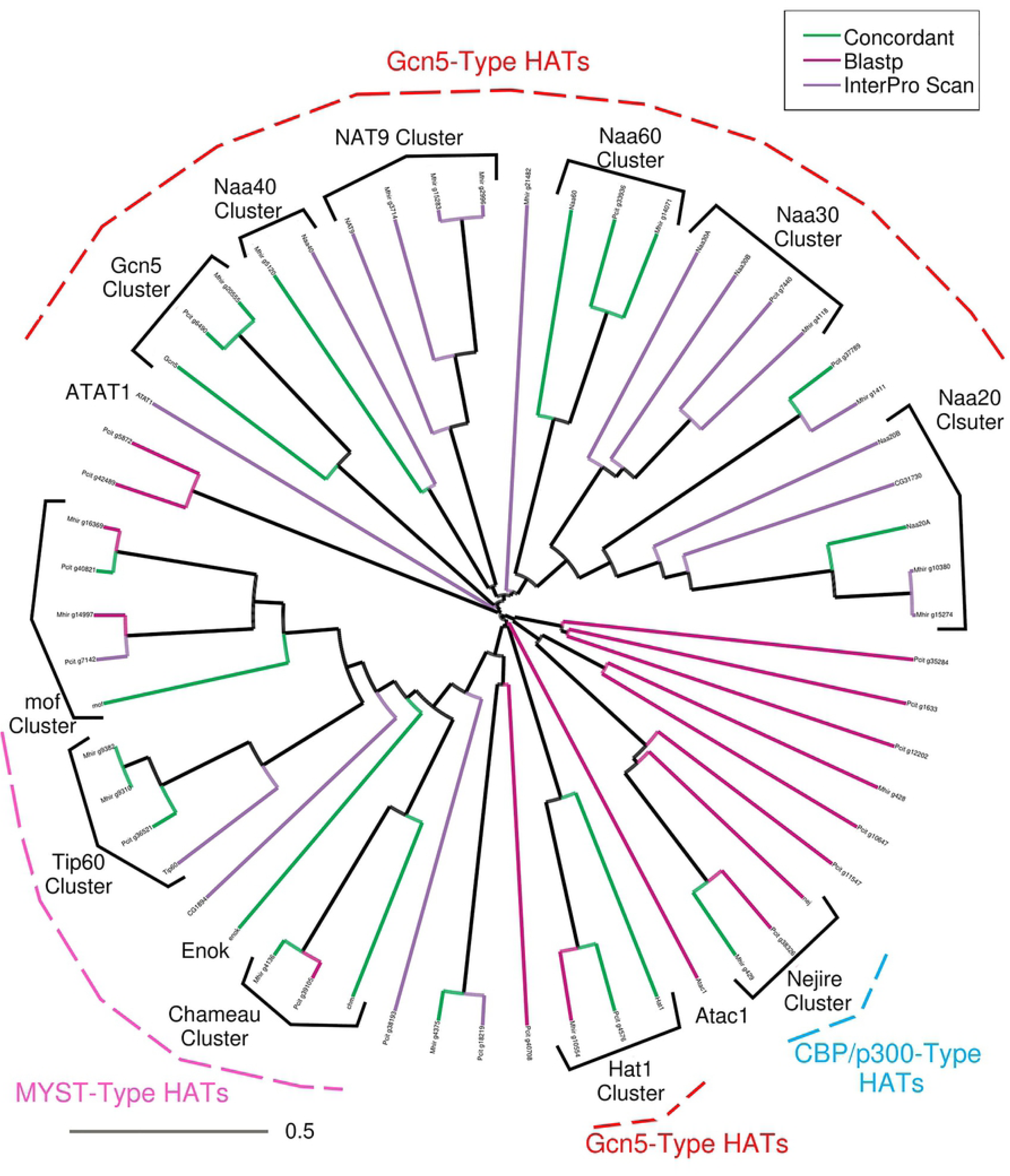
**Phylogenetic clustering of histone acetyltransferase (HATs).** The protein sequences of HATs of Mhir and Pcit, were aligned with those of *Dmel* as the reference sequence. The three classes (BLASTp only, InterProScan only and Concordant), are indicated by differently coloured lines, as given in the inset. The red, pink and blue lines outside the tree, indicate the clustering with a known Dmel proteins, confirming functional identity of Mhir and Pcit. The proteins from Mhir and Pcit have the suffixes Mhir and Pcit respectively, while the Dmel proteins are named according to the nomenclature in Uniprot.

The Gcn5-related HAT1 forms a distinct cluster with Mhir (g10554) and Pcit (g4576) proteins (Fig 11). The Dmel Gcn5 clusters also include Mhir (g20555) and Pcit (g6490) proteins. HAT1 is important for the *de novo* histone deposition and chromatin assembly which is in turn associated with HAT1-mediated cycle of H3 and H4 acetylation and deacetylation [100]. HAT1 also contributes to cellular tolerance to double strand breaks which are induced by replication blocks [101]. *Drosophila* Gcn5 catalyzes H3K9 and K14 acetylation and is a key player regulating metamorphosis and oogenesis [102]. The association of Gcn5 mutation with decondensation of the X-chromosome, similar to that found in case of mutations in Iswi and Nurf301, link X-chromosome condensation with histone acetylation [103]. Another well-known histone acetyltransferase, Nej harbours CBP/p300-type HAT domain and acetylates H3K18, H3K27 and H4K8. The products of Mhir g429 and Pcit g11547 and g38362 genes cluster with Dmel Nejire protein (Fig 11). A number of other Mhir and Pcit proteins cluster with lesser known class of Gcn5-related KATs called N(Alpha)-Acetyltransferases which are members of the NAT (N-terminal acetyltransferase) complexes (Fig 11)[104]. Not much is known about these GNAT domain-containing N(Alpha)-Acetyltransferases but their depletion alters global H3 and H4 acetylation levels [105]. HATs and HDACs cooperate to regulate allele-specific histone acetylation at the DMRs (Differentially Methylated Regions) [106]. Thus, the activating marks are important players in regulation of parental-origin-specific transcription.

The comparison of HAT genes between different species points towards expansion of this gene class in *A*pis(17), Dmel (16) and Mhir (16) with Pcit and Clec carrying only 10 and 7 HAT genes, respectively (S9 Table). Since, HATs participate in dosage compensation and imprinting, the large number of HAT genes in Mhir can be predicted to play a key role in dosage compensation associated with paternal chromosome-specific heterochromatinization via hyperacetylation of the maternal chromosomes, which is to be investigated. The various bootstrap phylogenetic trees for the HATs are given in S6A-G Fig.

### Erasers of epigenetic imprint

#### Histone demethylases

Histone demethylation confers reversibility to histone methylation which is important for selective activation and repression and also for meiotic memory as in genomic imprinting. These serve as erasers of epigenetic marks. There are two kinds of demethylases-those that demethylate through the activity of the amine oxidase domain, using FAD as a cofactor and those where JmjC domain participate in demethylation [107, 108]. The histone demethylase enzymes were mined based on the presence of HPD, JmjC domain in Mhir, Pcit and other insects.

The highest number of demethylases are present in *A. pisum*(26) followed byDmelandPcit having equal numbers of demethylases (13),Mhir has 12 and Clec has 16. A large number of demethylases are found in the BLASTp only class in mealybugs-Mhir (12) and Pcit (20) which need further validation (Fig9A,S9 Table). We used phylogenetic clustering with *Drosophila* HDMs and found multiple genes coding for the almost all demethylases in both Mhir and Pcit. Based on this clustering, the putative novel genes are identified as *JMJD4, Kdm4a/b, jarid2, JMJ14* and *HSPBAP1* (Fig12).

The amine oxidase family of demethylases consisting of LSD1/su(var)3-3 are found in Mhir (2 genes) and Pcit (3 genes; Fig12). Lsd1-mediates H3-K4 demethylation and in *Drosophila*, the mutants are sterile and defective in ovary development. The mutant alleles of Lsd1 in *Drosophila,* suppress position-effect variegation, suggesting a disruption of the balance between euchromatin and heterochromatin [109]. Both lid and dKdm2 target H3K4me3 and regulate transcription of essential developmental genes. They are required for different developmental processes but may have some redundant functions [110]. Mhir has 2 and Pcit has 1 copy of lid, whereas both Mhir and Pcit have 1 copy each of Kdm2. Kdm2 also demethylates H3K36 [111].

Kdm4a and Kdm4b genes, demethylate H3K9me3. In *Drosophila*, Kdm4A showed strong association at heterochromatin which led the authors to propose that Kdm4A is a structural component of heterochromatin [112]. Kdm4a regulates heterochromatin position-effect variegation (PEV), organization of repetitive DNA, and DNA repair [112]. Mhir has 1 copy of Kdm4 while Pcit has 2 copies (Fig 12). JHDM2 (JmjC domain-containing histone demethylation protein 2)/ Kdm3 is known to have demethylation activity at H3K9. Mhir contains 1 copy of each of Utx and Jarid2 (JmjC genes) while Pcit has two copies of Utx and one copy of Jarid2. In Dmel, mutation in three out of 13 Histone demethylases shows lethality [113].

**Fig 12.**
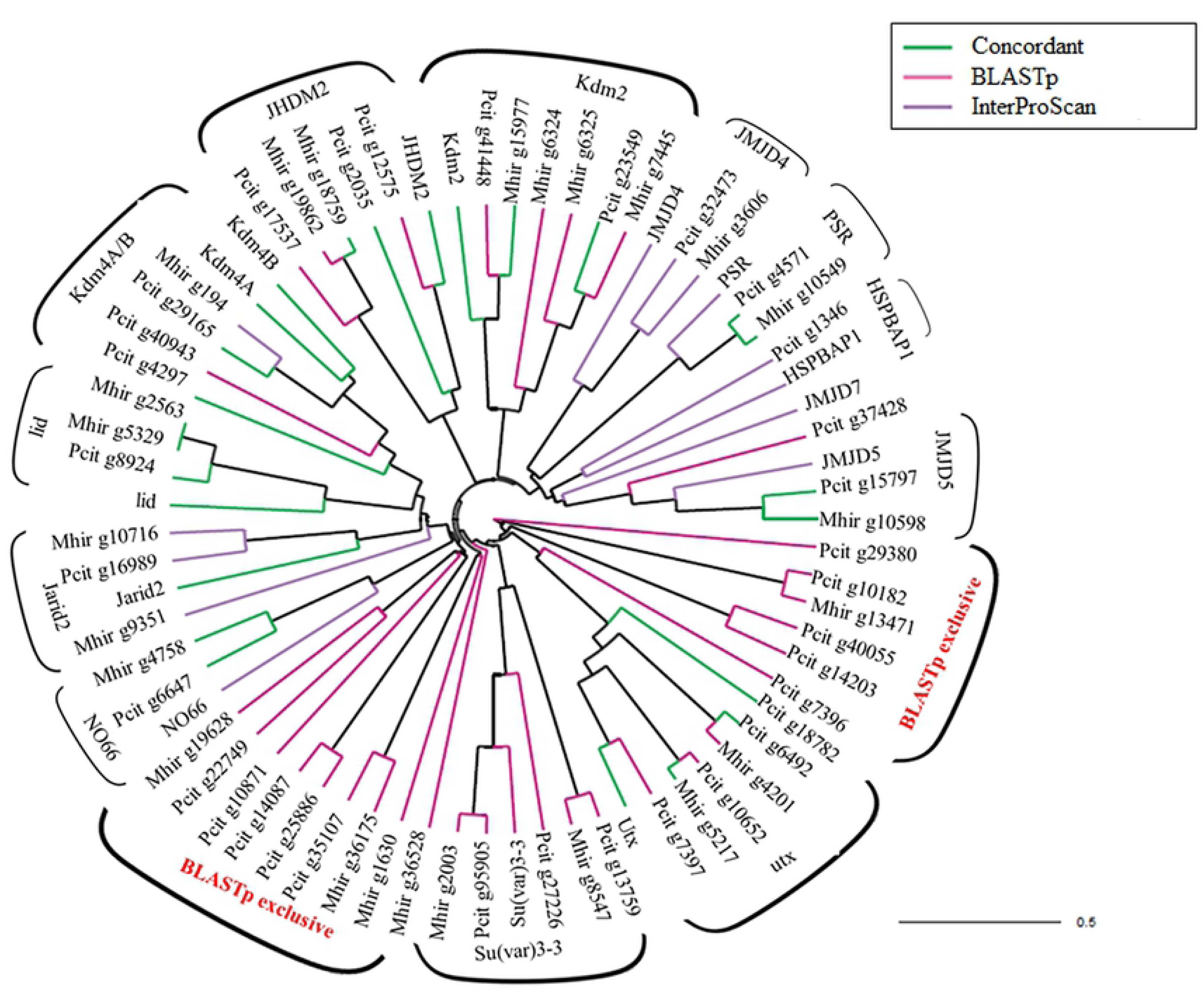
**Phylogenetic cluster for Histone demethylases (HDMs).** The protein sequences of HDMs of Mhir and Pcit, were aligned with those of Dmel used as the reference. The three classes (BLASTp only, InterProScan only and Concordant), are indicated by differently coloured lines, as given in the inset. The black lines outside the tree, indicate the proteins clustering with a known Dmel proteins, confirming identity of Mhir and Pcit genes. The proteins from Mhir and Pcit are indicated by suffixes, while the Dmel proteins are named according to the nomenclature in Uniprot. The BLASTp only members are indicated in red. Most of the putative novel demethylases cluster with JMJD4, HSPBAP1 and Jarid2 proteins of Drosophila.

JMJD7 demethylating arginine residues of histones H2, H3 and H4 in *Drosophila*[114] is absent in Mhirand Pcit. Both Mhir and Pcit have 1 copy each of JMJD4, JMJD5 and HSPBAP1/CG43320. JMJ14 encoding histone H3K4 demethylase is present in Mhir, but is absent in Dmel. NO66 specifically demethylating H3K4me and H3K36me of histone H3, plays a central role in histone code [115]. Both Mhir and Pcit have one copy of the gene (Fig12). The various bootstrapped trees for the histone demethylases are given in S7 Fig.

Since the demethylases act on specific residues of specific histones and absence of any report of genes to demethylate H3K79 and H4K20; it remains unclear how the histone demethylation is carried out at these sites in the Dmel, Mhir and Pcit genomes.The writers and erasers of histone methylation are detected in the mealybug genome conferring reversibility to histone methylation as an epigenetic signature.

#### Histone deacetylases

Histone deacetylases (HDACs), associated with transcriptional repression are categorized into four classes on the basis of DNA sequence similarity and function. Class I, II and IV enzymes are inhibited by trichostatin A (TSA) and known as the classical HDACs while class III members are NAD+-dependent proteins which are not inhibited by TSA and are known as Sirtuins. We have analysed the classical HDACs of Class I, II and IV in Mhir and Pcit using *D.melanogaster* as the reference.

HDAC1 plays a crucial role in imprinting and X-chromosome inactivation, via its interaction with NuRD chromatin remodelling and deacetylase complex [116, 117]. HDAC inhibition by TSA, leads to the loss of hypoacetylation associated with inactive X [118]. In *Drosophila*, HDAC1 and SU(VAR)3-9 co-operate to methylate pre-acetylated histones to bring about transcriptional silencing. The HDAC3 of Mhir (g3497) and Pcit (13687) cluster with HDAC3 of Dmel (Fig 13). The HDAC1 and HDAC3 clusters form a part of a larger cluster that includes other Mhir (g5630) and Pcit (g31540) proteins (Fig 13). HDAC3 binds to putative enhancers on the X-chromosome and promotes histone deacetylation upon *Xist* induction, promoting transcriptional silencing [119]. Global deacetylation by HDAC3 is a prerequisite for chromatin compaction during mitosis [120].HDAC3 via its association with linker histone H1.3 also regulates polar microtubule dynamics in mitosis thus controlling spindle formation and chromosome alignment [121]. The expression of this gene during development would be of interest in the mealybugs, where mono-polar spindle is formed during spermatogenesis [8].

**Fig 13.**
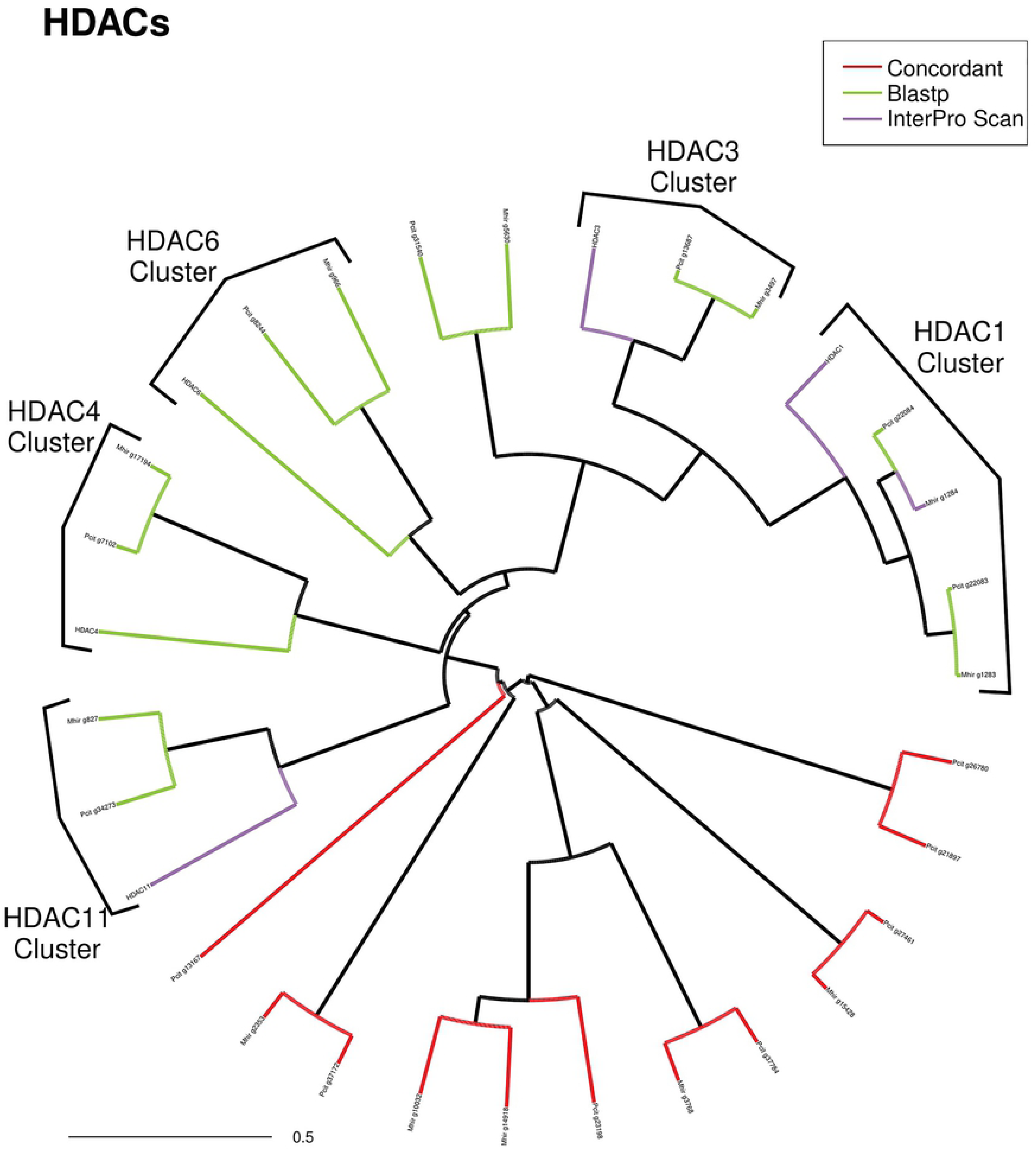
**Phylogenetic clustering of histone deacetylases (HDACs).** The protein sequences of HDACs of Mhir and Pcit, were aligned with those of *Dmel* as the reference sequence. The three classes (BLASTp only, InterProScan only and Concordant), are indicated by different coloured lines, as given in the inset. The black lines outside the tree, indicate the proteins clustering with a known Dmel proteins, confirming Mhir and Pcit functional identity. The proteins from Mhir and Pcit have the suffixes Mhir and Pcit respectively, while the Dmel proteins are named according to the nomenclature in Uniprot.

Mhir (g17194) and Pcit (g7102) proteins are orthologues of HDAC4, while Mhir (g966) and Pcit (g8244) cluster with HDAC6.Dmel HDAC11 clusters with Mhir (g827) and Pcit (g34275) proteins (Fig 13). HDAC11, the lone member of Class IV, is an unusual type, which is present in Mhir and Pcit, although it has not been identified in *A. pisum*[47]. Apart from its histone deacetylase activity, HDAC11 also acts as a fatty acyl-hydrolase [122]. All the clusters described above form a part of a supercluster that also includes Pcit proteins (g13167 and g37172) and Mhir protein (g2353). The bootstrapped trees for the HDACs indicate high conservation andpotential functional similarity (Fig14).

**Fig 14.**
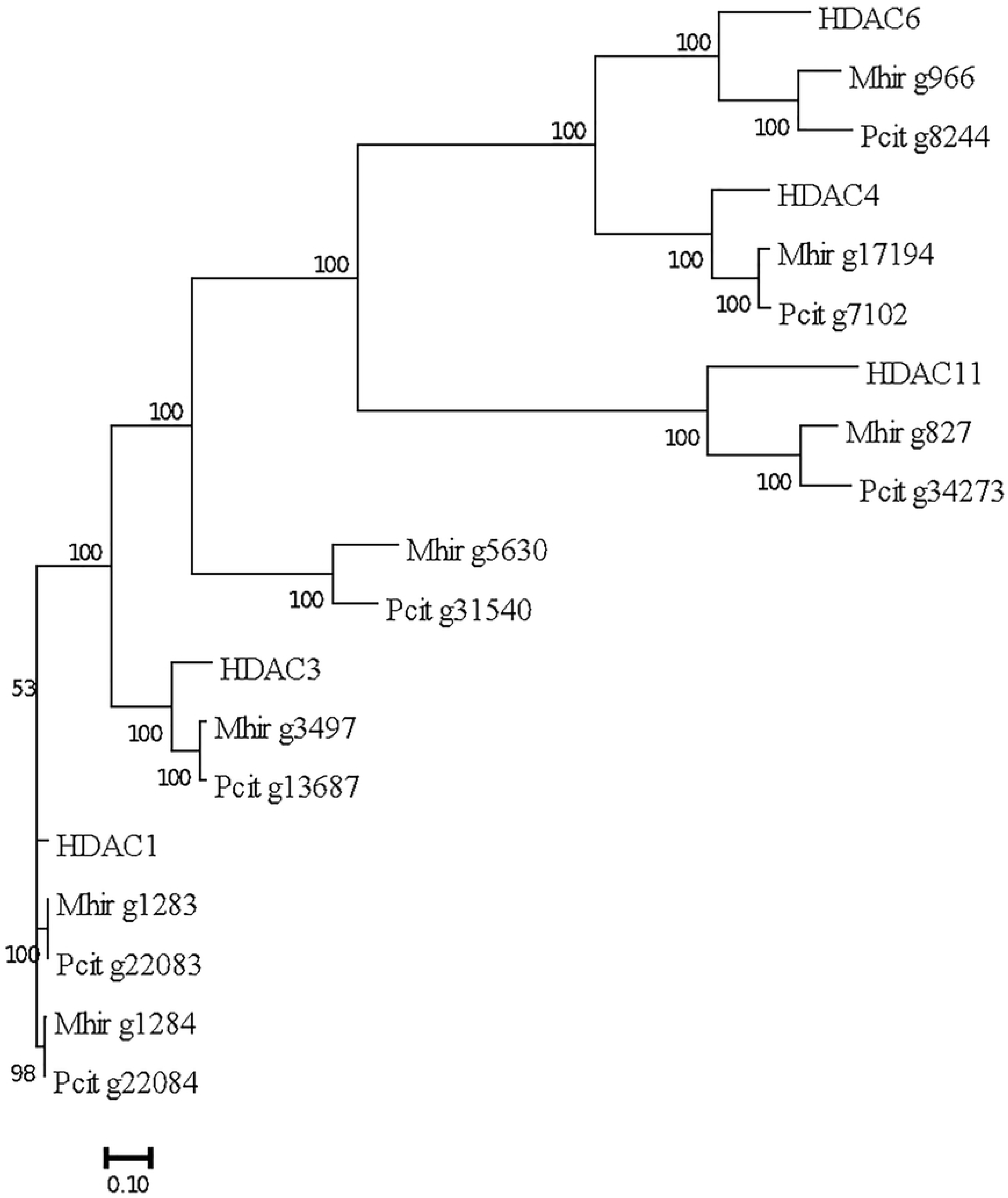
**The bootstrap phylogenetic tree for the histone deacetylases**. The numbers on the branches represent the bootstrap value assigned to each node.

In mealybugs, where the paternal genome is heterochromatized, a complete repertoire of classical HDACs working along with histone methyltransferases would be a pre-requisite. It is interesting to speculate that this repertoire for the heterochromatization of paternal chromosomes in mealybugs and X-chromosome inactivation in mammals are shared. HDACs appear as an expanded class in Mhir, Pcit, *A*pisand Clec with *A. pisum* harbouring maximum number of HDACs, when compared to Dmel. Evolutionary pressures such as genome expansions along with parahaploidy in mealybugs may have led to the expansion of chromatin modifier gene families, regulating high density chromatin packaging [123].

### Protein complexes mediating epigenetic modification in mealybugs

The epigenetic modifier proteins are recruited to their site of action through protein complexes that bring about chromatin accessibility and also mediate the recognition of the modified histones to translate the signal for either transcriptional activation or repression [124]. The genome of Mhir and Pcit contain almost all the catalytic activity required for epigenetic modifications of DNA and histones as well as those required for reversal of the modification. The target specificity of these enzymes is governed by protein complexes that recruit the writers and erasers. In light of these observations, the involvement of PcG-like (Polycomb) and TrxG/COMPASS-like complex inmealybugsis also necessary. These complexes are well conserved from yeast to mammals [125].Our current analysis aims to identify and compare the composition of Polycomb and Trithorax and Chromatin remodelling complexes in mealybugs, Pcit and Mhir. Using the well-studied complexes in *D.melanogaster* as the reference we analysed genes involved in these complexes in the Mhir and Pcit genome.

### Polycomb and Trithorax complexes

The Polycomb and Trithorax complexes are highly conserved across species although the DNA elements that recruit these complexes (PRE/TRE) are not conserved [126, 127]. We find that they are conserved in Mhir and Pcit with some of the being multi-copygenes. For example, in PRC2 complexes there are three Enhancer of zeste-like proteins in Mhir(g5597, g5598 and g1288) and two in Pcit (g13871 and g37973) indicating redundancy compared to Dmel(Fig 15). Similarly, in case of PRC1 complexes there are two RING finger protein 3-like proteins in Mhir (g14582 and g15883) as opposed to one dRING in Dmel PRC2 complex. There are cases where the Mhir and Pcit polycomb members are closer in sequence to human homologues than to the Dmel homologues; Mhir and Pcit YY1-like proteins (g4392 in Mhir and g1376 in Pcit) and RING finger protein-3 like (g14582 and g15883; Fig 15. Upon alignment, these proteins were found to share 60.5% and 61% similarity with human YY1 protein, respectively, whereas their similarity to Pho (*Drosophila* homologue for YY1) is 46.67% and 48.98%, respectively.

**Fig 15.**
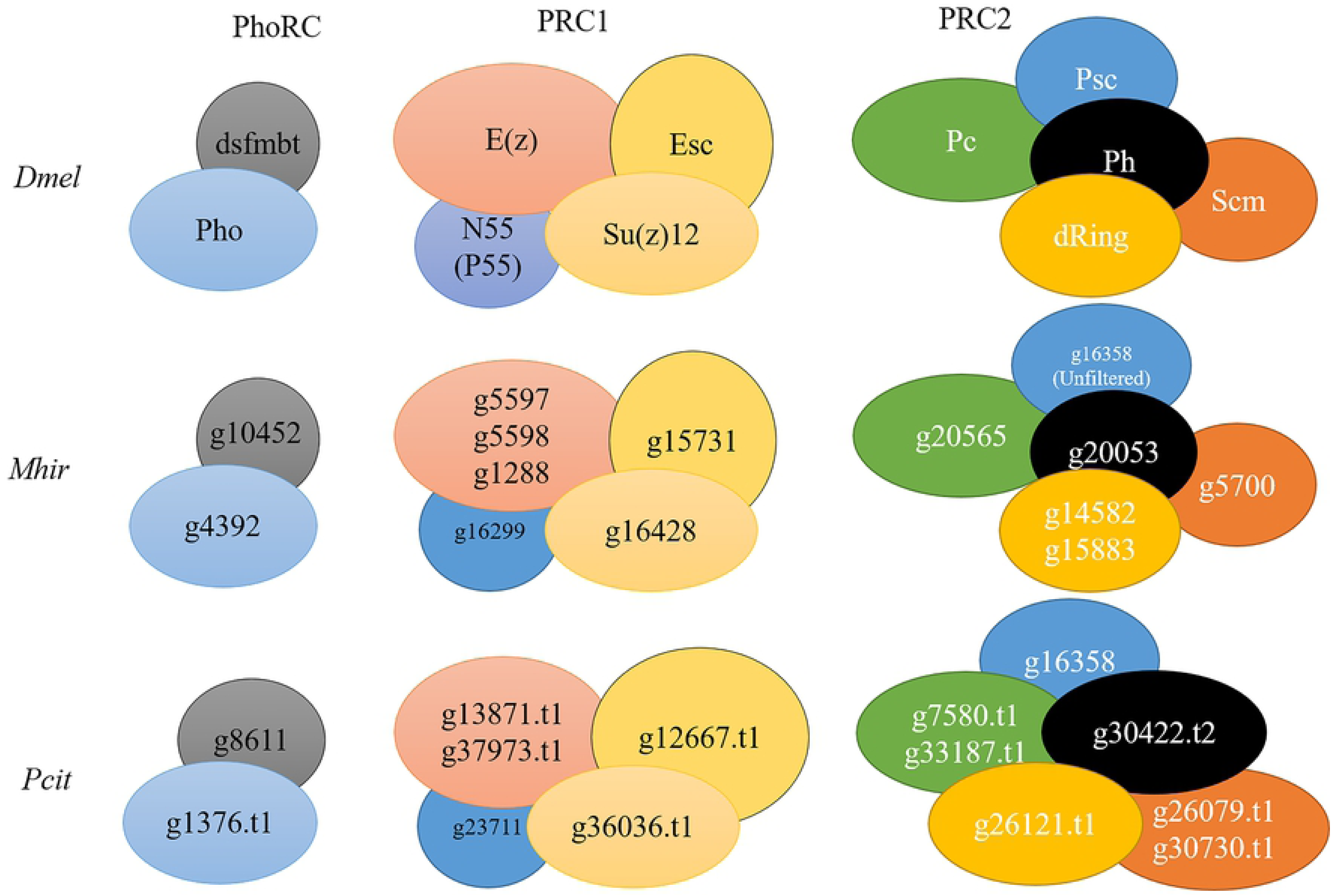
**Conservation of Polycomb Complexes between *D. melanogaster, M.hirsutus* and *P citri.*** The copy number of genes for some proteins of the in PRC 1 and 2 complex is higher in Mhir and Pcit. The Pho gene in Drosophila is the homologue of YY1 in mammals and the mealybug homolgue is closer to YY1 as discussed in the text. The colour coding is maintained to indicate the homologues in Dmel and the mealybugs.

The DNA binding domain of YY1, Pho, YY1-like protein of Mhir and Pcit are highly similar (≥94%) and they are different outside this domain. All the four proteins have four well conserved C2H2-type Zinc fingers and ZF2 and ZF3, which are essential for YY1-mediated transcription [128]. YY1 has a histidine cluster which is a nuclear speckle-directing sequence and is missing in Pho and the YY1-like proteins of Mhir and Pcit. The nuclear speckles are the centre of RNA synthesis and processing. The histidine clusters appeared after the duplication event associated with the vertebrate evolution [129]. The HAT/HDAC interacting domain varies between YY1 and other proteins, even though the REPO domain, that participates in PcG recruitment to DNA and is essential for PcG mediated repression, is well conserved between YY1, g4392 and g1367 [130]. The spacer sequence in YY1 and Mhir/Pcit YY1-like proteins share greater similarity and differs significantly from that of Pho protein. The spacer regions act as accessory regions to transactivation function and the deletion of the spacer at the C-terminal end, perturbs the DNA binding and transactivation activity [131].

RING finger proteins are members of the PRC1 complex, which bring about E3-ubiquitin ligase activity. The zinc finger associated with the RING finger domain is highly conserved between *Drosophila* dRing/Sce, Mhir (g14582 and g15883) and Pcit (g26121) RING finger 3- like protein, even though the rest of the sequence is not conserved. Another interesting feature is the complete absence of the RAWUL (Ring Finger/WD40 association ubiquitin-like) domain at the C-terminal in the RING finger 3-like proteins of Mhir and Pcit. The RAWUL domain contains a ubiquitin fold and is important for interaction with the Cbx members of the PRC1 complex [132]. The Dmel E(z) protein and E(z)-like proteins of Mhir (g5597, g5598 and g1288) and Pcit (g13871 and g37973) have high sequence similarity, except for the partial deletion in the SANT-Myb domain. The SANT-Myb domain interacts with the nucleosome and the inter-nucleosomal DNA. It would be interesting to see how the partial deletion of the SANT-Myb domain affects the activity of Mhir E(z) and whether it is compensated by another accessory protein harbouring the SANT-Myb domain.

The Polycomb complex is a writer as Ezh2 catalyses H3K27 di/trimethylation, leading to condensed chromatin and gene silencing. These complexes are recruited to the chromatin with the help of transcription factors (example Pho/ YY1 binding to GCCAT/ACCAT) [133] or short- and long-non-coding RNAs [134, 135]. The presence of YY1 like proteins in the mealybug genome is significant in this context. The analysis of the non-coding RNA is underway.

The writer for signalling activation is the Trithorax Complex, also known as the COMPASS complex which brings about H3K4 di/trimethylation leading to an open chromatin state and transcriptional activation (Fig 16). The trithorax complexes are also well conserved in Dmel, Pcit and Mhir. But there are examples where the Mhir or Pcit homologues were not identified for example, dNcoA6 of Trithorax-related dCOMPASS-like complex is missing in Mhir and Pcit, complexes (Fig 16). Ash2 is missing in the Pcit in Trithorax dCOMPASS-like complexes. The absence of the genes in both Pcit and Mhir genome may indicate true absence in mealybug genome, while absence in only one of the two, like absence of Sh2, suggests sequence gaps.

**Fig 16.**
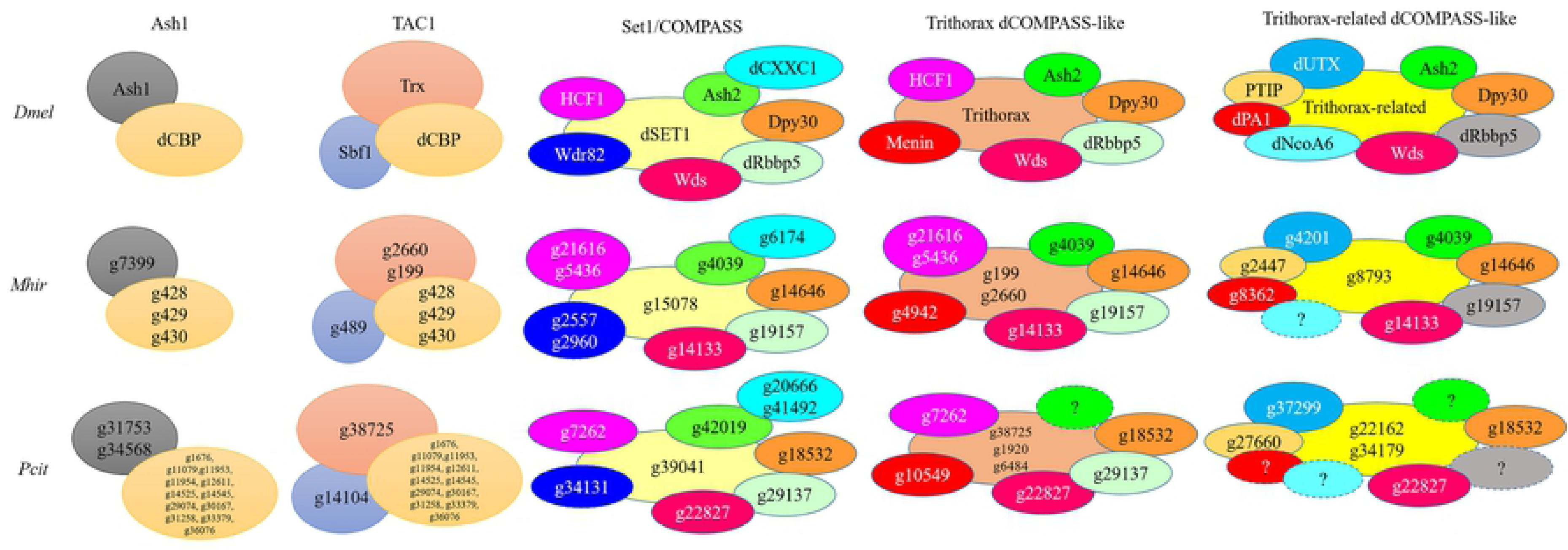
**Conservation of Trithorax Complexes between *D. melanogaster, M.hirsutus* and *P citri*.** Multiple copies of some of the homologues is indicated. The missing proteins in the mealybugs is indicated (?). The colour code corresponds to the specific protein in Dmel and gene(s) in mealybugs.

There are multiple genes which are dCBP-like in Mhir and Pcit suggesting functional redundancy or tissue specific expression. The *Drosophila* dCBP (CREB-binding protein) is also known as Nejire that harbours CBP/p300-type HAT domain and acetylates H3K18, H3K27 and H4K8. Nejire-mediated H3K18 and H3K27 acetylation controls male sterility in *Drosophila* [136]. Thus, its role in sexual dimorphism in mealybugs can be speculated. Thus, the current analysis points towards the presence of a complete repertoire of Polycomb and Trithorax complex members in Mhir and Pcit with few exceptions.

### Chromatin remodelling complexes- the readers of epigenetic signals

The structure of chromatin and its dynamics is essential for active transcription as well as for the selective compaction of the chromatin in transcriptionally silent regions. In epigenetic regulation, the readers (chromatin remodelers) recognize the histone modification as one of the signals and remodel nucleosomes thereby facilitating the compaction/expansion of the nucleosomal arrays.

The high-priority domains utilized to identify the chromatin remodeling genes (CRM) are the Helicase ATP binding, SNF2 N-terminal, Helicase C and the P-loop NTPase domains (S2 Fig). The InterProScan and the BLASTp analysis of Mhir genome led to the identification of 30 chromatin remodelling proteins, including 2 putative novel chromatin remodelers (S11 Table). We carried out a comparative analysis of all the genes in Mhir with other insect genomes (S11 Table). *D.melanogaster* has well defined candidates for CRM genes along with some predictedchromatinremodelers that contain the HPD (S11 Table). It is known that the *A. pisum* genome is duplicated for the epigenetic modifiers including the chromatin remodeling proteins [137]. Pcit genome has several genes that are annotated as chromatin remodelers only in BLASTp analysis.

The chromatin remodelers are classified into 4 families, namely SWI/SNF, ISWI, CHD and INO80. Proteins of these four chromatin remodeling complexes identified in Mhir and Pcit compared with *Drosophila* complexes are shown in Fig17, S8 A-C Fig. BLASTp analysis identified a complete repertoire of these proteins in Mhir genome indicating functional conservation. A number of proteins in these complexes harbour additional domains that recognize various histone modifications and bring about nucleosome sliding, histone variant exchange and/or nucleosome ejection [138, 139].

The core proteins of the various chromatin remodeling complexes present in Mhir and Pcit are shown in Fig17 (SWI/SNF) and S8Fig. It is observed that there are multiple copies of genes for several proteins within the complex, as in the case of BRM protein in BAP and PBAP complexes of the SWI/SNF family (Fig 17) and Domino of SWR1 complex (S8AFig). Similarly, there are 3 NURF38 coding genes in Mhir and 2 in Pcit (S8B Fig). These proteins are similar to the Dmel proteins in having the HPD, but may have additional domains. The Mi-2 protein has a potential DNA binding domain [140]. It is possible that the expression of these genes is either tissue specific or developmental-stage specific. The increase in copy number also suggests the importance of the function of the gene and hence, the evolution of redundancy.

**Fig 17.**
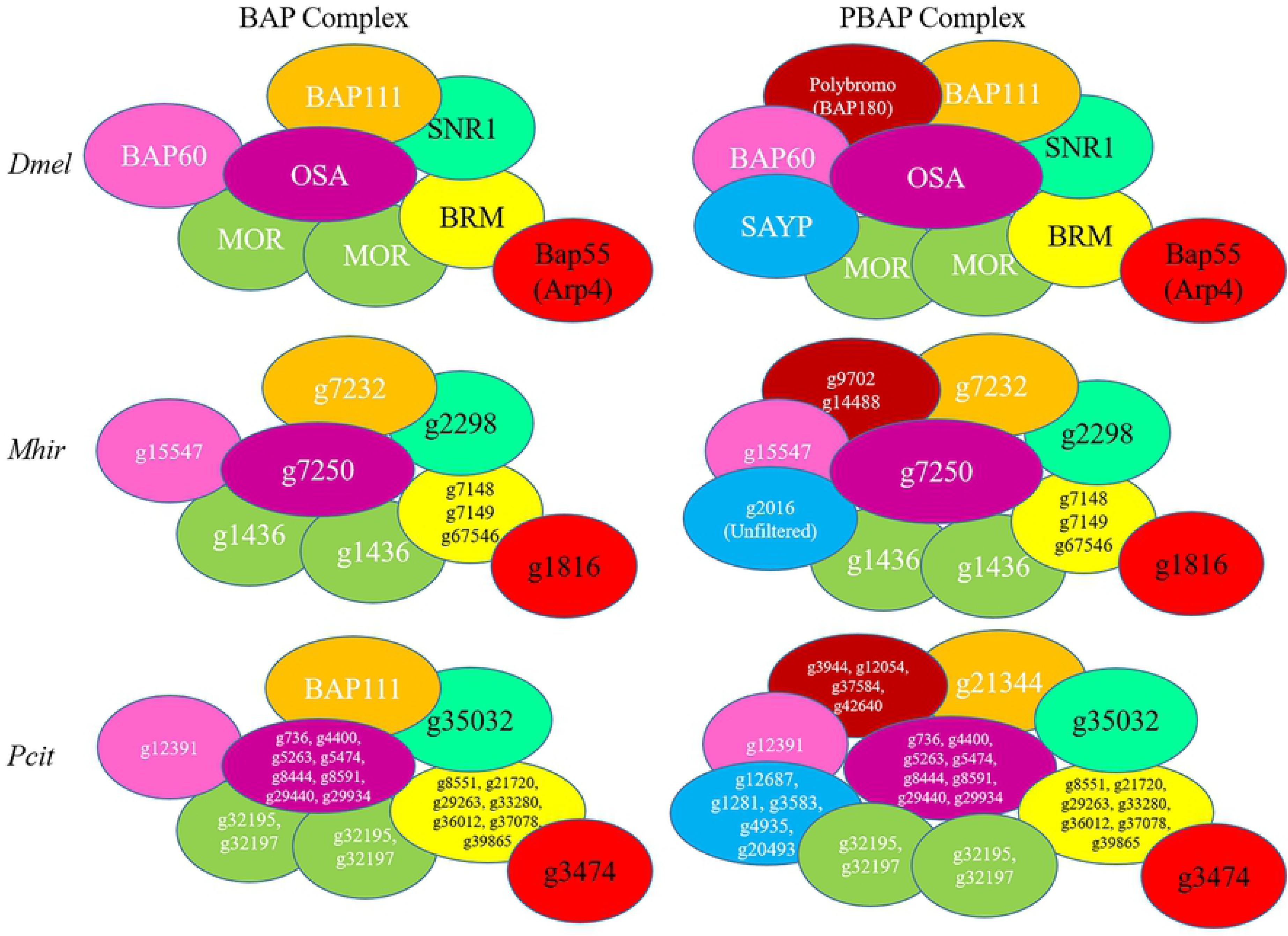
**Conservation of SWI/SNF complexes between *D. melanogaster, M.hirsutus* and *P citri***. These complexes are complete and some homologues occur in multiple copies. The colour code corresponds to the specific protein in Dmel and gene(s) in mealybugs.

Apart from the core complex, which is highly conserved across species, the composition of the accessory proteins varies in tissue and developmental stage-specific manner. The recruitment of the various complexes to their site of activity is generally through recognition of the histone modification of the site. A summary of selected examples of histone modifications recognized by various chromatin remodeling complexes is given (S12 Table). The writers for all the histone modifications that mediate the recruitment of the chromatin remodelling complex are coded in the Mhir and Pcit genome, as discussed earlier. This shows that the mechanisms of epigenetic regulation known in other systems can be functional in the mealybug system. It is interesting that some of their gene structure is similar to the human genes rather than that of Drosophila.

### Transcriptome analysis for differential gene expression

Transcriptome sequencing of adult females and males was performed and the data statistics is given in Table 2. Analysis of the transcriptome indicated concordance between the biological replicates (Fig18). We analyzed the transcriptome of endosymbionts (mapped genes contributed by endosymbionts) and *M.hirsutus* nuclear genome. It was found that on an average endosymbionts (*Candidatus Tremblaya princeps* and *Doolittlea endobia)* have higherexpression in females than males (Fig 18). This correlates with the earlier observation that the endosymbiont load is reduced in the non-feeding adult males of mealybugs. In two mealybug species (*Planococcus kraunhiae* and *Ps.comstacki*), the analysis of the dynamics of infection of both beta and gamma proteobacteria indicated comparable level in both males and females in the early stages of development, while it was detected only in adult females. Similarly in *Planococcus lilacinus* 16s rRNA was found only in adult females [141, 142]. The elimination of endosymbionts from adult males could be due to their reduced metabolism as they do not feed, they mate with several females and die after a few days [143]. Unlike the endosymbiont transcripts, host gene transcripts showed no significant difference between adult male and female mealybugs (Fig 18).

**Fig 18.**
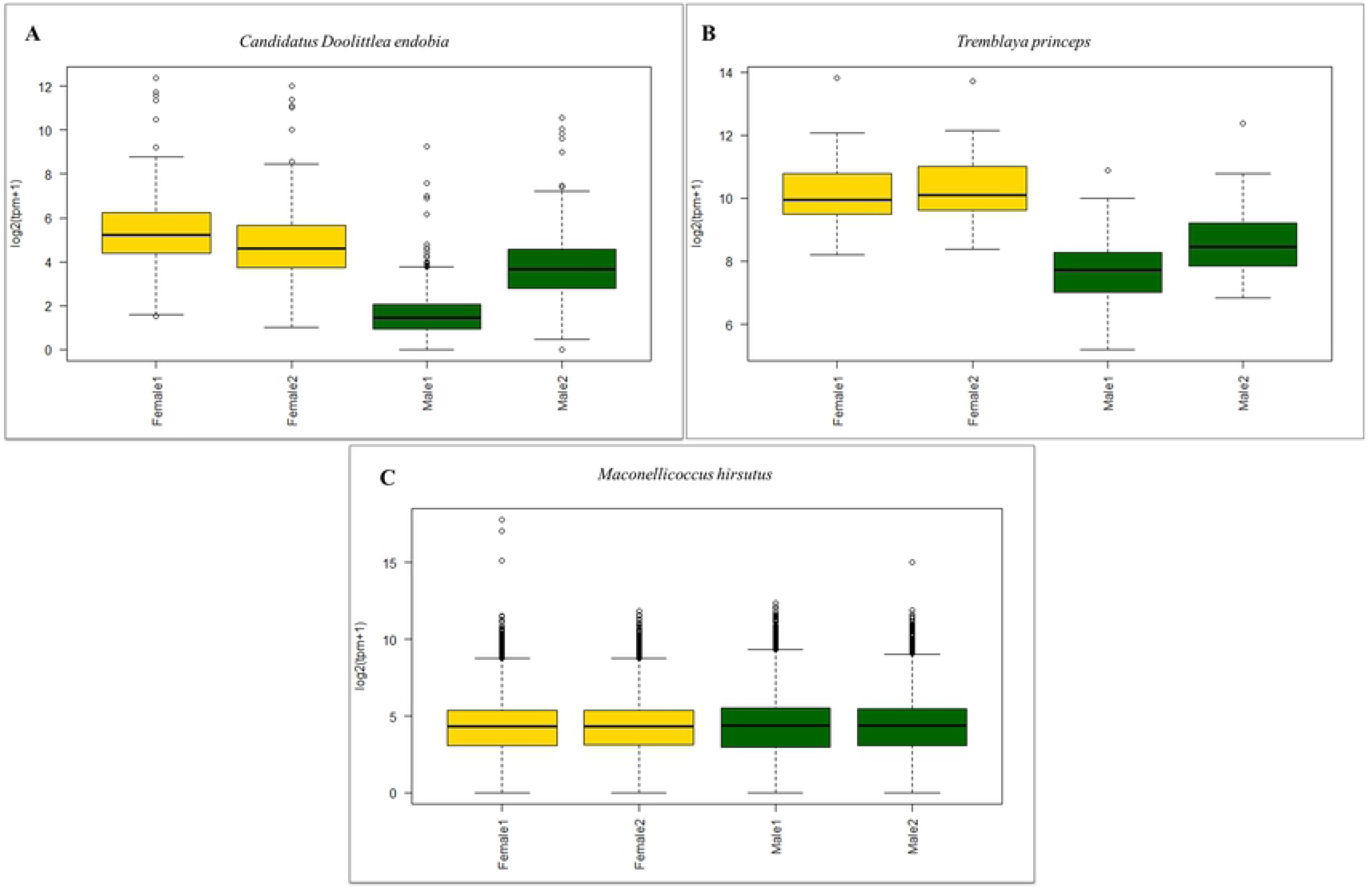
**The distribution of transcripts from the mealybug genome and the two nested endosymbionts in male and female mealybugs.** A- *Candidatus Tremblaya princeps*, B-*Doolittlea endobia,* C- *M. hirsutus* in the transcriptome of male and female mealybugs. A significant difference in transcript abundance from both the endosymbionts is observed in males and females, but not for transcripts from the mealybug genome.

**Table 2:** Read and alignment statistics of RNA-sequence data.

| Sample | Paired<br>endRaw<br>reads | Filtered<br>reads | % of data<br>remaining after<br>applying filters | % of data aligned to<br>dataset1 [Annotated<br>gene set] |
| --- | --- | --- | --- | --- |
| Male Replicate 1 | 38255062 | 3,61,52,236 | 95 | 49.46578131 |
| Male Replicate 2 | 44675238 | 4,26,70,610 | 96 | 47.19257587 |
| Female Replicate 1 | 26842830 | 2,49,41,567 | 93 | 48.58446544 |
| Female Replicate 2 | 40545144 | 3,81,84,609 | 94 | 55.38986663 |

The number of paired end raw and filtered reads as well as percent alignment

Differential gene expression (DGE) analysis was performed using the Kallisto-Sleuth pipeline and 1183 genes were found to be differentially expressed in males and females after applying multiple filters (S9 Fig). Hierarchical clustering of the differentially expressed (DE) shows similarity between the biological replicates and variation in the expression of genes among males and females (Fig 19). Out of the 1183 genes, 652 genes have higher expression in males and 531 genes show higher expression in females, these will be referred to as male enriched and female enriched genes hereafter in this manuscript.

**Fig 19.**
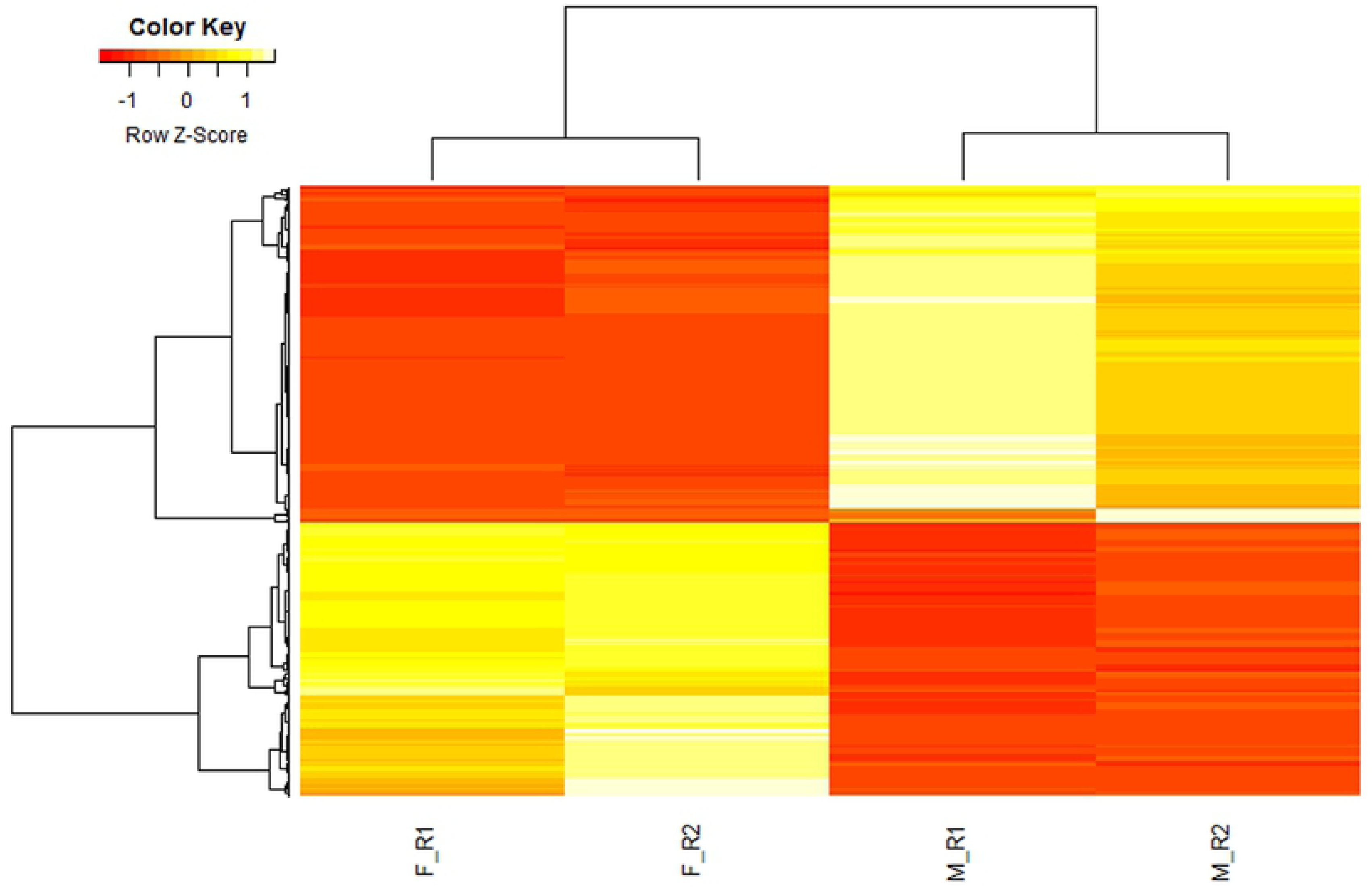
**Heatmap depicting hierarchal clustering of differentially Expressed Genes, (DEGs)) in male and female replicates.** Based on q-value <0.05 and log_2_FC > 1, 1183 genes were identified as differentiaaly expressed genes. F_R1 and F_R2 are two female replicates and M_R1 and M_R2 are two male replicates. Expression scale is defined by color key (top left).

We used a combination of approaches to find functionally enriched pathways and processes in males and females. We found genes related to metabolic and oxidative phosphorylation pathway enriched in males, while genes of ribosome biogenesis pathway are enriched in females (S10 Fig). Further we removed the genes related to oxidative phosphorylation and ribosomesl from genes overexpressed in males and females respectively and performed GO annotation. Biological processes show enrichment of genes related to “translation” and “response to oxidative stress” specifically present in females while genes related to “cytoskeletal organization” and “cellular protein modification” were present in males (S11Fig). In addition, we performed manual curation of all DE genes (S12 Fig, Fig20&21) which indicated enrichment of genes involved in metabolism, signal transduction, transporters sensory transduction and insecticide resistance in both males and females. Biological functions specifically enriched in males include cytoskeletal organization, cuticle development, protein ubiquitination and autophagy; while in females, genes involved in translation, ribosomes, RNA processing and wax biosynthesis were enriched. The significance of these observations was considered in the light of life-cycle and the sexual dimorphism of the mealybugs in terms of their size and morphology.

**Fig 20.**
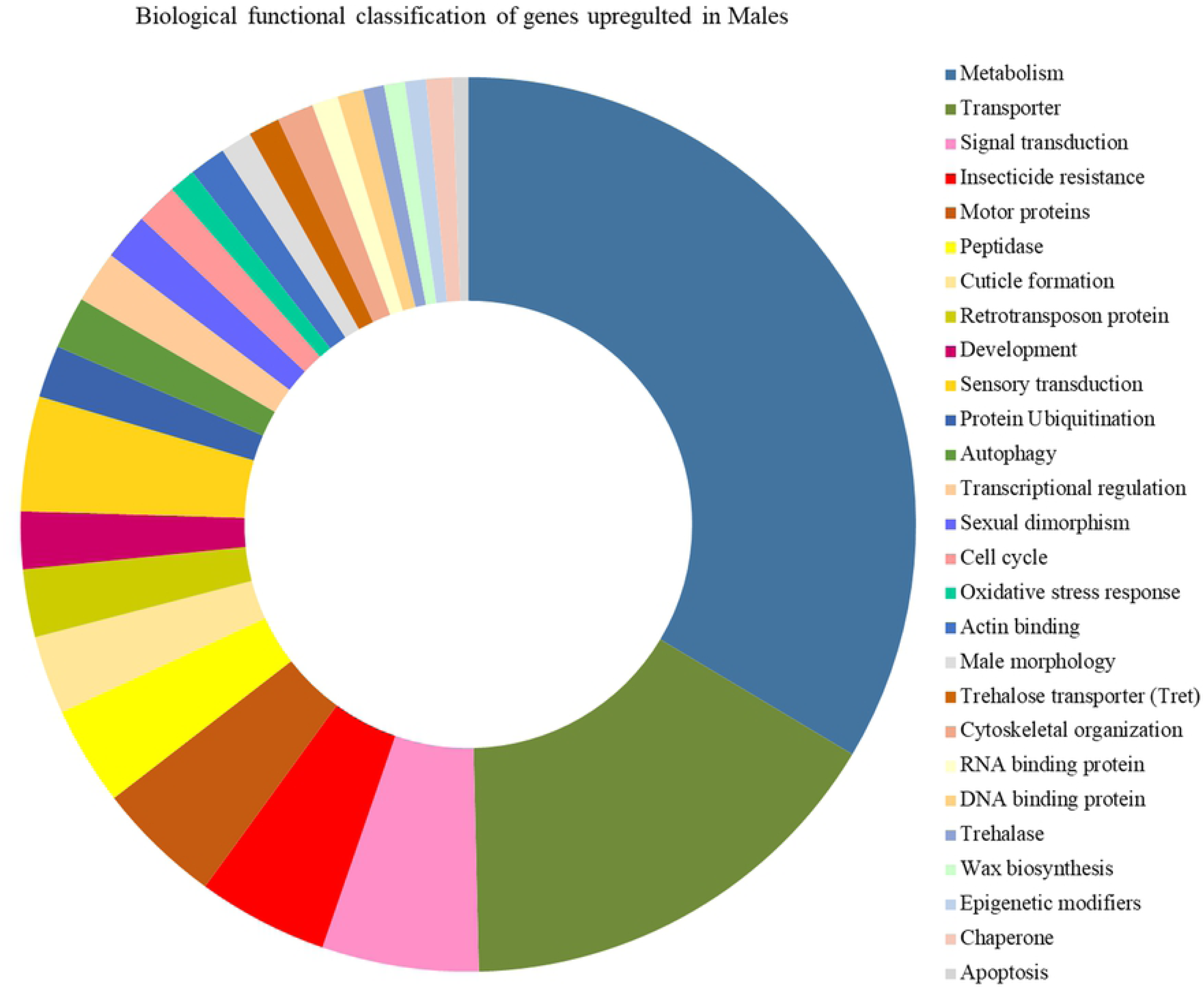
**Biological function based classification of genes with male enriched expression in *M. hirsutus* from transcriptome data.** The most enriched classes are metabolism and transporter class.

**Fig 21.**
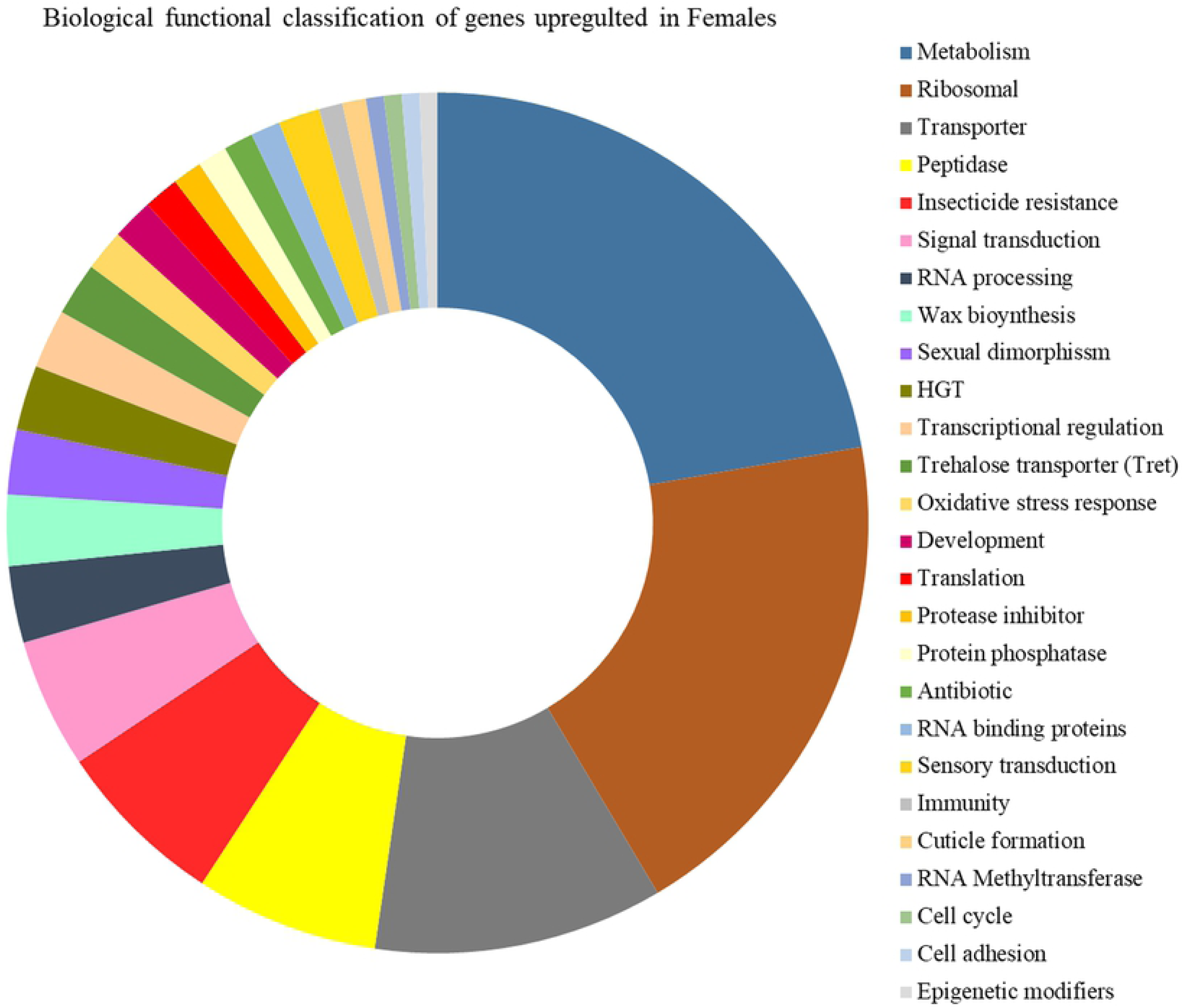
**Biological function based classification of genes with enriched expression in females.** Transcripts from genes for metabolism and ribosomal functions are most abundant.

In both males and females, the most prominent pathways with increased expression are those for metabolism (Males: 177; Females: 100) though the nature of metabolic pathways varied. In males oxidative phosphorylation genes (70), lipid metabolism (16), fat metabolism (16) and TCA cycle genes (14) are over-represented. In females a relatively high number of DE genes are from carbohydrate (21), lipid (15) and fat metabolism(15). This enrichment can be attributed to high energy requirements for flight in adult males for which lipids and fats are utilized, and oxidative phosphorylation pathway is involved in energy production. On the other hand females are constantly feeding to provide for growth and development of eggs thus would require genes for carbohydrate metabolism.

Manual annotation of DE genes highlighted gene classes involved in metamorphosis, insect flight and chitin synthesis over-expressed in males. These include four copies of trehalase genes which play a critical role in metamorphosis, insect flight and chitin synthesis [144]. Motor proteins like flightin, myosin, tropomyosin, paramyosin, troponin C and alpha-actinin which form part of the insect flight muscle [145] also show elevated expression in male mealybugs.

The enrichment of transcripts in males that correlate with sexual dimorphism are the following: flightin gene, specifically found in flight muscles and essential for flight, with fold change of 8.8, takeout (g1473) involved in courtship behaviour and autophagy pathway which is induced by starvation and is one of the mechanisms adopted for elimination of endosymbionts in cereal weevil Sitophilus [146]. Higher expression of flightin gene is also observed in winged morphs in *Aphis gossypii*[147, 148].

Reciprocally in females, increased expression of genes correlating with oocyte development is seen. The genes involved in ribosome function, RNA processing, RNA binding, transcriptional regulation and translation also can be attributed to deposition of maternal transcripts and proteins during oocyte development. One of the ribosomal proteins RPL12, interacts with trithorax and polycomb complexes and deregulates heat-response and ribosomal protein genes [149]. Mhir has one copy of the RPL12 gene (g17594) and its transcription is higher in females than in males. Apart from this, genes for wax biosynthesis are enriched, correlating with secretion of wax filaments by adult females to form ovisac [150].

As mentioned earlier, several horizontally transferred genes (HGTs) identified in mealybug involved in protein degradation, Vitamin B and amino acid metabolism showed increased expression in females. This enrichment of HGTs and endosymbiont specific transcripts in females correlated with their lifespan and growth, in contrast to males.

We divided the DE genes into 3 categories based on fold change (FC): FC 1.5-2.9, FC 3.0-4.9 and FC 5.0-10. There were more genes (30 genes) from males in the high fold change category (FC5.0-10) of differential expression compared to females (4 genes). The GO classification of these genes indicated that the four genes overexpressed in females are for carbohydrate metabolism. On the other hand the genes highly expressed in males belonged to multiple functional categories (Fig 22) and consist of a variety of genes relating to energy production which in turn can be related to sexual dimorphism including flight,courtship and mating which are male specific attributes.

**Fig 22.**
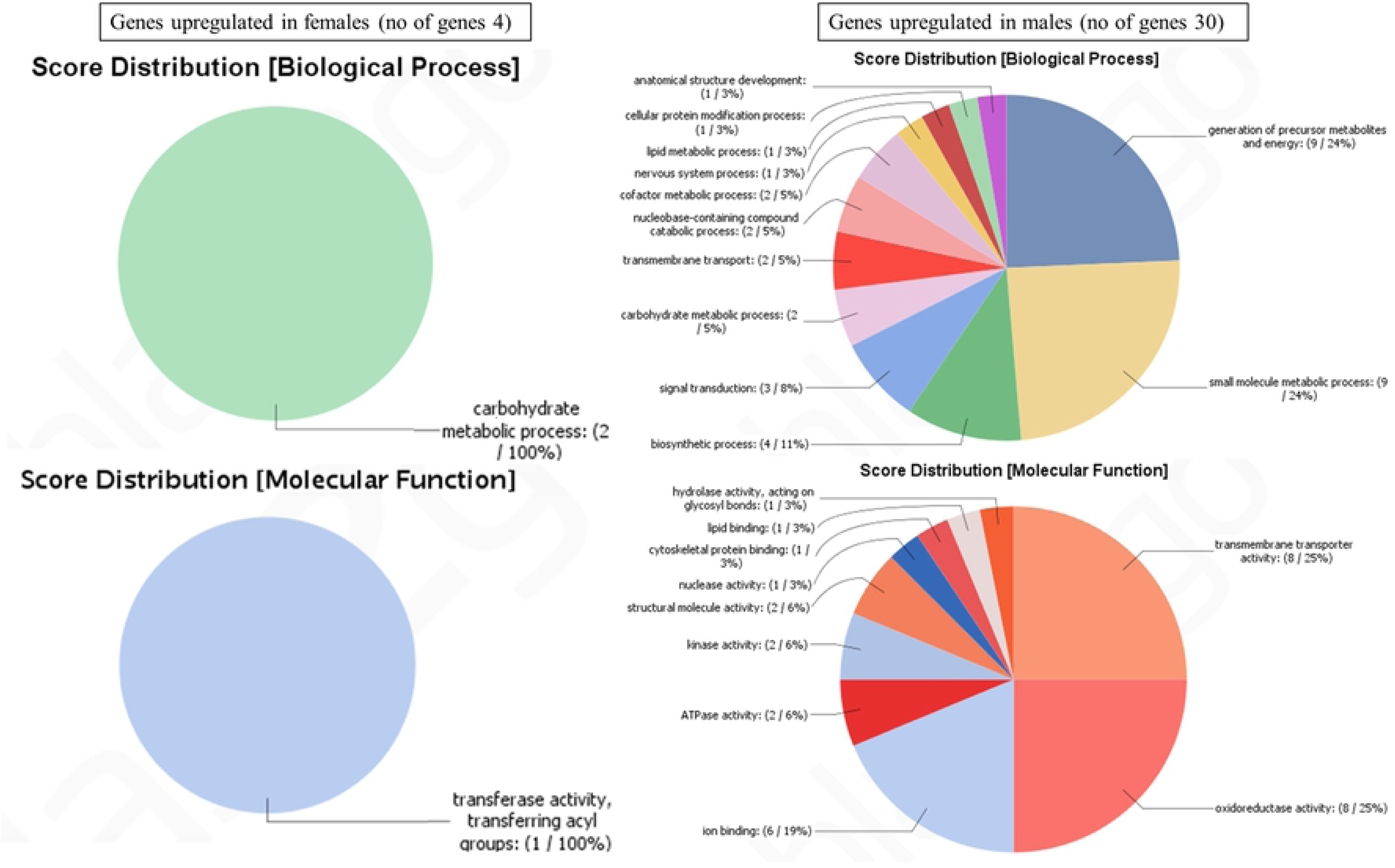
**BLAST2GO derived GO classification of genes having 5-10 fold difference in gene expression between females and males.** The different GO categories represented include biological process and molecular function terms enriched in genes up-regulated in females (left) and males (right). There are larger number of genes highly over-expressed in males and they represent multiple functional class.

### Expression of epigenetic regulators

All the epigenetic regulatory genes in Mhir are expressed and a few genes showdifferential expression (S13Table). *SMYDA-5*, *SMYDA-4* and *SDS3* have increased expression in males, *SMYDA-5* and *SMYDA-4* share homology with human SMYD3 protein, that can methylate H3K4me3 and histone H4 at lysine 5 (H4K5) and lysine 20 (H4K20) [151]. In Drosophila, they are associated with histone deacetylase binding activity and negative regulation of gene expression.

SMYD5, *SMYDA-5* and nucleoplasmin show increased expression in females. *SMYD5* over-expressed in females, brings about H4K20me3 modification [152]. This modification is associated with DNA damage repair, chromatin compaction and heterochromatin formation. The establishment of heterochromatin depends on the recruitment of H4K20 histone methyltransferase by H3K9me3 bound HP1 [153]. This H3K9me3-HP1-H4K20me3 pathway is shown to regulate facultative heterochromatization in *Planococcus citri*, with both repressive methylations localizing to the heterochromatin in male mealybug nuclei [15]. SDS3 forms an integral component of Sin3 histone deacetylase corepressor complex playing a critical role in its integrity and is essential for its deacetylase activity [154].

Apart from the DE genes, other epigenetic modifier genes identified in Mhir genome, are expressed in both adult males and females. We analyzed the expression of these genes and identified genes that showed differences in expression between males and females based on TPM values (S13 Fig). Several genes such as *SUV420H1* (*Su(var)4-20*, g12552), *PR-Set7* (g14923), *SUV39H2* (*Su(var)3-9*, g6525), *E(z)* (g5598) and *Jarid2* (g10716) associated with repressive histone methylation show increased expression in males along with *JMJ14*(g9351) involved in demethylation of active histone mark H3K4me3. However, these did not cross the threshold criteria of fold change (FC) ≥ 1. Overall the expression of all epigenetic modifier genes is moderate to low as seen in terms of relative TPM values (S14 Table). This low expression in adult males and females may be an indication of the maintenance state of already established epigenetic marks. It is important to investigate the transcription of these genes at different developmental stages to correlate their activity with genomic imprinting mechanisms.

In X chromosome inactivation in female mammals repressive histone marks like methylation and also the removal activating marks by histone deacetylation are essential [116, 117, 119]. It remains to be investigated if SMYD proteins along with H4K20 methyltransferase like Su(var)4-20 are part of the imprinting machinery in the mealybugs, for maintenance if not for initiation.

### X-chromosome inactivation, a comparable paradigm

The differential regulation of homologous chromosomes in mammals and the mealybugs are well known examples of facultative heterochromatization in two evolutionarily distant species. The process of X inactivation in mammals is random as opposed to that in the mealybugs, though the similarities in late replication, transcriptional repression, and the establishment of inactivation occurring around a similar developmental timeline, suggest evolutionary conservation of the process [4, 155]. The mechanisms for the selection of the homologue for inactivation in diploid cells differ in the two systems. However, the mechanisms of formation of heterochromatin and its maintenance through mitosis may bear similarities, beyond the epigenetic modifiers of histone and DNA. In this context, we examined proteins involved in facultative heterochromatization of chromosomes for differential regulation in mammals and the mealybugs.

Based on the available literature on X chromosome inactivation, we considered the protein factors that interact with the long non-coding RNA as assembly factors to establish inactivation. The presence of homologues of genes coding for these proteins in the mealybug genome are identified (S15 Table and Fig 23). We have considered the molecular process leading to chromosome condensation and transcriptional inactivation in three different phases, however this does not correlate to the temporal sequence of these events.

**Fig 23.**
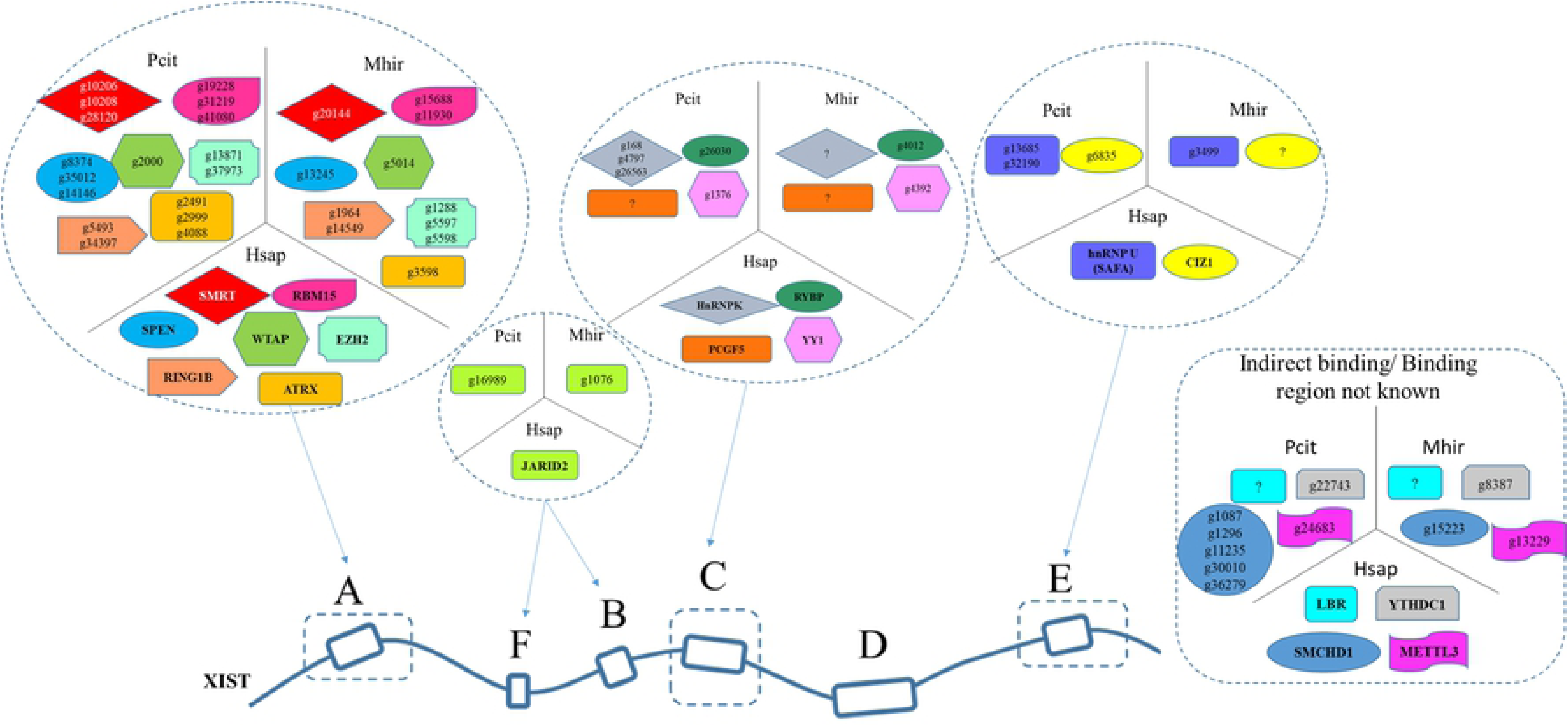
**Proteins for facultative heterochromatization shared between mammals and the mealybugs.** The homolgues of almost all proteins that interact with XIST RNA are conserved in Pcit and Mhir. The copy number of the homolgues for certain proteins are higher in Mhir and Pcit. Hsap is for Humans.

I: Removal of activation marks is an essential step to achieve inactivation as seen in different systems, including the mealybugs [95]. The SPEN protein (SMRT/HDAC1-associated repressor protein) carries out HDAC mediated histone deacetylation leading to transcriptional repression. Mhir has a single copy of *SPEN* while 4 copies are present in Pcit. The domain analysis revealed that among the four genes only one gene of Pcit (g35012) contains SPOC domain essential for *SPEN* function.

RBM15 is for recruitment of METTL3(RNA m6A methyltransferase) to Xist RNA in a WTAP-dependent manner for mRNA modification. Allthe genes for these functions are found in Mhir and Pcit with *WTAP* being present in single copy and *RBM15* in multiple copies in both the genomes.

II. Localization of Xist to inactive X: The proteins implicated in this process are nuclear matrix proteins SAF-A, CIZ1 and transcription factor YY1 ([156, 157]). *SAF-A* is present in both Mhir and Pcit, while *CIZI* is identified in Pcit, but not in Mhir. The transcription regulator YY1 is the mammalian homologue of the pleiohomeotic (Pho) of Drosophila. The mealybug homologue is similar to YY1 rather than Pho as discussed earlier.

III. Addition of repressive marks: The PRC1 and PRC2 complexes contribute to heterochromatin assembly on Xi through H3K27me3 and H2AK119 ubiquitination (H2AK119Ub; [157]). EZH2 of the PRC2 complex deposits the repressive histone marks and JARID2 helps in recruitment of the complex to the inactive X chromosome. The chromatin remodeller, ATRX2 serves as a bridging factor to reinforce PRC2 recruitment to inactive X [156, 157]. The genes for these proteins are present in both Mhir and Pcit genomes, except genes for *PCF2* (in Mhir) and *HnRNPK* (in Mhir and Pcit). It remains to be seen if there are other proteins which can substitute for these. The other important proteins not detected in mealybugs are lamin B receptor and macroH2A. The list of the proteins discussed here and their status in mealybug genome is given in S15 Table.

The detection of most of the protein factors interacting with Xist RNA in the mealybug genome suggests evolutionary conservation of mechanism of facultative heterochromatin. The major player, the Xist-like long non-coding RNA remains to be identified in the mealybugs. On the other hand, the presence of proteins that interact with Xist is a robust indicator of the presence of such a RNA or atleast the conservation of consensus sequence motifs either in a long non-coding RNA or in DNA itself. One of the important differences between X inactivation and the inactivation of paternal chromosomes in the mealybug is the well supported evidence suggesting the presence of multiple centres of inactivation in mealybugs [16, 158]. It is to be noted that the multiple copies of the genes essential for inactivation may be due to a large proportion of the genome (50%) being subjected to inactivation in the mealybugs.

In summary the analysis of mealybug genome considering it as a model for genomic imprinting reflects the conservation of the molecular players of whole chromosome inactivation on one hand, and on the other variations as in DNA methyltransferases, may reflect novel processes. The genome analysis reflects the basis of the unique biology of mealybugs in terms of radiation resistance, DNA repair and other features. The genome and the transcriptome described here provide a resource for further work.

## Materials and methods

### Establishing mealybug culture

*Maconellicoccus hirsutus* (Mhir) was collected from an infected custard apple. The individual gravid females were cultured on sprouting potatoes. The identification of the species was carried out based on the cuticular features [159]. A colony of sexually reproducing Mhir was established and is maintained since 2012. The colony is maintained on pumpkins at 24-26^0^ C in dark glass chambers with fine mesh on one of the four sides. A pool of embryos was collected by placing gravid females on crumpled parafilm, laid out on agar plates with pumpkin extract.

### Genomes for comparative analysis

The genome data of following insects were used for comparative analysis: *Drosophila melanogaster* (Diptera)as a well-studied model system, the pea aphid *Acyrthosiphon pisum*, the triatomid bug *Rhodnius prolixus and Cimex lectularius* belong to the order hemiptera to which the Mhir and Pcit belong, the silkworm *Bombyx mori* (Lepidoptera), which is plant feeder. The availability of well annotated genome data and the variation in habitat were also considered for choice of these systems for comparative analysis.

### Genome sequencing and annotation

The genomic DNA was extracted from the pool of embryos using phenol-chloroform method. The genomic DNA (gDNA) library for sequencing was prepared according to the instructions by the manufacturer for sequencing on HiSeq2000 platform (Illumina, USA) and the Ion Torrent PGM platform (Life Technologies), PacBio sequencing was outsourced to Genotypic Technology Pvt. Ltd., India. Illumina paired-end data was filtered using Trimmomatic (version 0.35;[160]) and merged to form longer super-reads using MaSuRCA.3.2.1 [161]. The filtered Illumina reads were used to error correct PacBio reads using PBcR pipeline of Celera assembler (version 8.3). Error corrected PacBio reads having length ≥500bp were selected and finally the RunCA module was used to generate the final assembly using a combination of Illumina super-reads and error-corrected PacBio reads. Pilon was further used to correct the assembly. Ion torrent PGM contigs were constructed using CLC Genomics Workbench 9 to validate the assembly.

*Ab initio* gene prediction was done using RNASeq based genome annotation tool BRAKER_v1.9 [162] which combines prediction from the tools GeneMark and Augustus to identify the final set of genes/proteins in Mhir genome. TheRNASeq reads from the Mhir embryos were aligned using STAR and were provided as an input to the software. To train the gene prediction model for Augustus, parameters were automatically optimised from transcriptomics assisted GeneMark predicted genes in the prior step.

BLASTp (version BLAST 2.4.0; [163]) was used for annotating the function of the predicted protein sequence set using NR database as the reference. Maximum five hits were considered for every query. The high confidence hits with E-value <10^-3^, percent identity (p.ident) ≥ 30, query coverage (qcovs) ≥50, and successful alignment with RNA reads were considered for analysis. Taxonomy was retrieved by configuring NCBI taxonomy database (taxdb) into the BLAST analysis. Manual curation was carried out to validate the annotation. InterProScan tool (version 5.19-58.0; [164]) was used for identification of the domains/protein signatures. Protein domains with e-value <0.001 were selected for analysis. In addition, the genome of Mhir, Pcit and the others were analysed using InterProScan for the presence of different domains to predict the function of hypothetical proteins of selected classes. The genes that did not have any domain and covered less than 30% of reference sequence in BLAST were considered truncated and not considered for the downstream analyses.

### Validation of assembly

The assembly of the genome was validated by PCR using tiling primers, followed by Sanger sequencing. The primers were designed considering gene architecture as in assembled scaffold and the sequence obtained was compared with the same scaffold. The scaffold containing histone genes was selected for this analysis. The primer sequences are given in S16 Table. Long PCRs (5-7 kb amplicons) were also carried out for validation of these scaffolds.

### Detecting Horizontal Gene Transfers (HGTs) in mealybug genome

The taxonomic classification of EggNOG [165] was used to find the genes of bacterial origin in the insect genome. The genes from endosymbiont scaffolds were discarded and ensured that HGTs have genes of Arthropoda origin in the same scaffold/contig. Also, the truncated proteins which did not qualify the QC applied in BLASTp analysis (percent identity ≥ 30% and query coverage ≥ 50%) were also discarded. Only those proteins were selected that contain at least one domain. HGTs were also validated using PCR.

### Detection of expansion and contraction of gene classes in M. hirsutus

For evaluating the gene class expansion and contraction, the proteomes of six insects- *P.citri, A. pisum, R.prolixus, C. lectularius* (hemipterans), *Drosophila melanogaster* (dipteran) and *Bombyx mori* (lepidopteran) were retrieved from NCBI (https://www.cbi.nlm.nih.gov) and used for all the gene model comparison with Mhir.

OrthoFinder (version 2.2.6) was used to compute and compare the gene numbers in orthologous clusters from seven proteomes of arthropod species *M. hirsutus, P. citri, A. pisum, R. prolixus, C. lectularius, D. melanogaster and B. mori*. The gene numbers in different orthogroups were compared to assess contraction and specific expansion of gene families or orthologous clusters in *M. hirsutus* genome. The gene counts of Mhir and Pcit were compared with the other five species. For avoiding bias, the endosymbiont specific orthogroups were removed from the analysis. To find the genes and gene families that were expanded, contracted or specifically present in Mhir and Pcit, the following criteria was applied-

i. To be considered as expansion, the maximum value obtained for gene counts of any of the five species should be less than the gene counts either in Mhir or Pcit.
ii. To find mealybug specific gene expansion, we included additional criteria for non-zero values in Mhir and Pcit while gene count of zero in the other five species,
iii. To identify contracted gene classes, the criteria is that the gene counts in both Mhir and Pcit must be zero while it should be non-zero value in all other species considered for comparison.

The functional annotation of the expanded, specific and contracted classes was carried out using EggNOG.

### Functional annotation of epigenetic regulators

**(a) Curation of epigenetic modifiers by identification of high priority domains:**

Manual curation was carried out, in addition to BLASTp, to validate the annotation using InterProScan (version 5.19-58.0) which contains a compilation of domains/protein signatures of genes based on the presence of functional domains. The domains were identified in Mhir, Pcit, Dmel, ApisandClec for comparative analysis. The domains with e-value <0.001 were selected for analysis. For example, to curate genes for epigenetic writers, readers and erasers, high priority domains were selected by computing the frequency of functional domains in genes of each class in *Drosophila melanogaster* while an in-house perl script was used to fetch genes containing the high priority domains [64]. After manual curation, these genes were divided into three sets; genes exclusive to BLASTp or InterProScan and those with concordant functional assignment in both BLASTp and InterProScan. BLASTp exclusive sets were analysed to further filter inaccurately identified members, those lacking functional domains [64].

### Phylogenetic analysis

For phylogenetic analysis of all the classes, InterProScan exclusive set was aligned with BLASTp exclusive and concordant dataset using MAFFT (version 7.3.94) with specific parameter L-INS-I, leaving gappy regions for better accuracy. Post-alignment, the phylogenetic trees were drawn using MAFFT and interactive visualization was obtained by Rainbow tree [166].

The InterProScan exclusive and the concordant datasets were aligned by MAFFT and MEGA7 software tool was used to create bootstrapped trees.

### Transcriptome sequencing and analysis

In the mealybugs, the male and female instars are indistinguishable upto the second instar stage, but can be easily identified as the third instars. The uninseminated female mealybugs were collected from cultures where the male development was inhibited by pyriproxyfen, an analogue of ecdysone [167]. The mealybug culture was seeded and maintained on sprouting potatoes, which was sprayed with 0.01ppm of pyriproxyfen for 21days and after this period, the culture was examined to confirm the absence of males and the females collected were the source of RNA. From a separate culture the winged males were collected and the absence of females was confirmed before extraction of RNA by TRIzol method [168].

The library prepared using the Illumina TruSeq stranded RNA library preparation kit following the manufacturer’s instructions. Ribo-Zero was used to deplete ribosomal RNA followed by fragmentation and priming for cDNA synthesis. The cDNA first strand was synthesized followed by the second strand synthesis. Adenylation of the 3’ ends was performed to prevent them from ligating to one another during the adapter ligation process. Following PCR enrichment, the concentration was estimated using Bioanalyser 2100 (Agilent Technologies) and sequenced on Illumina HiSeq 2500 producing 100X100-nt paired-end reads. Two biological samples were sequenced in duplicate. The RNASeq reads from adult male and female samples were filtered using Trimmomatic-0.36 and aligned to the annotated gene set from Mhir genome using Kallisto v0.43 with strand specific alignment parameters.

We sequenced the RNA from a mixture of male and female embryos at various stages of development. This data was used only to align with the genome assembly but not for any expression analysis as we did not have biological replicates.

### Differential Expression analysis

To analyse differential expression (DE) between male and female samples, Kallisto-Sleuth (v0.43.1 [169, 170]) pipeline was used. The raw RNA sequence reads from adult male and female samples were filtered using Trimmomatic-0.36. Kallisto was used for aligning filtered reads on annotated Mhir gene set. Sleuth was used to perform differential gene expression analysis. The genes with the threshold of 5% FDR (q-value < 0.05) were considered for the analysis. Additional filter criteria were also used to remove false positives. All the truncated genes as well as those genes that did not pass the QC criteria of BLAST annotation along with the endosymbiont genes were excluded. Only the genes having an expression of ≥1 TPM (Transcripts Per Kilobase Million) in at least two samples were selected for further analysis.

For computation of Log fold changes (LogFc), mean TPM values for both male and female samples was calculated, followed by the log2 scaled ratio of these average TPMs.For dealing with the null expression values, an arbitrary value of 1 was added to TPM values and finally, the genes with LogFc >1were selected for analysis. If the average expression in males is higher than females, it is termed as “Male enriched” and vice-versa. The complete expression data of all thegenes with their TPM values and fold change is given in S17 Table.

For hierarchical clustering of genes and samples, TPM values were transformed into z-scores using R and hclust function was used for performing hierarchical clustering and heatmap.2 was used for visualisation. A combination of approaches used for pathway enrichment and functional classification of differentially expressed genes include STRING, KEGG and BLAST2GO analysis. STRING (v 11.0) protein network with *Drosophila* as reference (in some cases the human data used) was drawn followed by KEGG pathway annotation to obtain pathways enriched in genes having higher expression in males and females respectively. For identification of the function of other DE genes in males and females, GO classification after STRING annotation was performed, after excluding the oxidative phosphorylation (from male-enriched) and ribosomal protein (from female-enriched) genes. Since several genes could not be annotated with STRING using *Drosophila* as the reference, BLAST2GO [171] was also used. Furthermore, manual curation for functional annotation of all DE genes we carried out as well (S12 Fig).

## Acknowledgements

We acknowledge DBT Bioinformatics facility at ACBR. Research fellowship from the following sources are acknowledged; PG-Dr. Shyama Prasad Mukherjee Fellowship from CSIR, SK-Senior Research Fellowship from UGC, JM-Research Associate fellowship from SERB No.60(0102)/12/EMR-II, AN acknowledges University Grants Commission for DS Kothari PDF fellowship.

## Supplementary figures

**S1Fig Circadian rhythm pathway genes contracted in Mhir genome**. The genes are compared with those of *Drosophila melanogaster* (Dmel) and Humans (Hsap). The core genes of the pathway that are absent in mealybugs are shown in grey box with broken line, *Tim* (*Timeout*) a paralog of *Timeless,* is present in the mealybugs which may compensate for the lack of Timeout present in Mhir.

**S2Fig High priority domain identification.** The frequency of the occurrence of the domains in HATs (histone acetyltransferase), HDACs (histone deacetylase) and CRMs (Chromatin remodeling) proteins in Drosophila is shown as an example. The frequency of occurrence of each domain is plotted as percentage on the Y-axis, # indicates high priority domain.

**S3Fig. Correlation of bootstrap-phylogenetic tree with domain architecture of proteins from Mhir and Pcit with Dmel as the reference.** A-Phylogenetic tree for E(z), B-comparison of domain architecture of E(z) gene from Drosophila, Mhir and Pcit. Mhir_g18633 is the E(z) protein identified only by BLASTp and is deficient in the high priority domains. C- Phylogenetic tree for the histone methyltransferase trr. D- domain architecture. Mhir_g13137 and Mhir_g20142 are the trr proteins identified by only BLASTp and are deficient in the high priority domains.

**S4Fig.The bootstrap- phylogenetic clustering of the histone methyltransferases** A- Set1, trr and trx, B- G9a and Su(var)3-9, C- Ash1 and Set2, D- PR-Set7, E- Gpp, and Ash2 F- HMT4-20. The numbers on the branches represent the bootstrap value assigned to each node. The genes identified only by BLASTp are excluded.

**S5Fig. The phylogenetic comparison of the arginine methyltransferase of Mhir and Pcit with that of Dmel.** A) Bootstrapped tree. B) Alignment of the proteins of Mhir and Pcit with that of Dme as the reference, (i) Art9 vs Art7 (ii) Art8 vs Art6 (iii) Art1 is shown with different colours with their alignment scores mentioned in the inset table. The table shown represents the proteins of the organisms- Dmel, Mhir and Pcit along with their length and their percent identity with the respective Dmel proteins. The line diagram from BLASTp were modified.

**S6Fig. The bootstrap-phylogenetic tree for the histone acetyltransferases**. The sequence conservation is reflected. A- mof, B- Gcn5, C- CBP/p300, D-Chm, enok, E-Tip60, F- Naa and G-NAT9.

**S7Fig. The bootstrap-phylogenetic tree for the histone demethylases**. A-Kdm4A, Kdm4B, lid and Jarid2 B-JMJD7, JMJD5, JHDM2 and JMJD4, C-Utx, D-HSPBAP1 and PSR. The genes identified only by BLASTp are excluded.

**S8Fig. Conservation of Complexes between *D. melanogaster, M.hirsutus* and *P citri.***The colour code corresponds to the specific protein in Dmel and gene(s) in mealybugs. A- INO80 family B-ISWI Complexes, C- CHD Complexes

S9Fig. Workflow followed and filters applied for transcriptome data analysis and identification of differential gene expression in male and female mealybugs.

**S10Fig. KEGG pathway enrichment among genes with higher expression** in A- females and B- males. Enriched GO terms identified in C- male upregulated genes after removal of oxidative phosphorylation related genes and D- female upregulated genes after removal of ribosomal genes. [I don’t know where this belongs]

**S11Fig. BLAST2GO derived GO classification of differentially expressed(DE) genes** in females (269) and males (319). The DE genes set after excluding oxidative phosphorylation related gene set from male and ribosomal genes from female were subjected to BLAST2GO.The Biological processes represented in females(A) and males (B) and the molecular processes in female(C) and male (D) are shown.

S12Fig. Workflow followed for manual curation of differentially expressed genes in males and females to classify them into different biological function categories.

**S13Fig. The difference in expression of epigenetic modifiers between males and females. The data is plotted interms of TPM values.**.

## Legends for Supplementary Tables

**S1 Table: Summary of the DNA sequencing data for Ion Torrent platform.** The sequence data before and after filtering the reads for quality is shown

**S2 Table: Summary of the genes identified as horizontally transferred genes.** The expression status is based on transcriptome data. *Considered if ≥ 10 reads mapped to the gene; **TPM> 0 values obtained from Kallisto sleuth pipeline.

**S3 Table: Summary of gene orthogroups expanded, specific and contracted in mealybugs *M. hirsutus* and *P. citri* genomes.** The number of genes present in each insect species in each orthogroup is given.

**S4 Table: Gene Orthogroups of Carboxylesterases.** Highlighted (yellow) Orthogroups represented in Spider Plots in Figure 4 in the manuscript.

**S5 Table: Gene Orthogroups of Cytochrome P450.** Highlighted (yellow) Orthogroups represented in Spider Plots in Figure 4 in the manuscript.

S6 Table: Orthogroups of genes missing in mealybug genome categorized according to their function.

**S7 Table: Copy number of DNA methyltransferases, demethylases and methyl CpG binding proteins present in mealybugs and other insect species.** * 1 bacterial origin, # identified as N6 DNA demethylase by BLASTp but clustering in human TET protein supercluster.

S8 Table: Identity matrix showing percentage similarity based on multiple sequence alignment of DNA demethylase proteins of different insect species from group 1 and 2 of Cluster C of phylogenetic tree.

**S9 Table: Comparative analysis of the number of genes coding for various histone modifiers of Mhir and Pcit genome with other insect species.** The numbers of genes for the writers and erasers are compared. The genes identified by BLASTp only did not have the high priority domains (HPD), those identified by InterProScan only had HPD, but were marked as hypothetical/unknown in BLASTp. Concordant classes were annotated by BLASTp and InterProScan. The genes identified by InterProScan only are the potential novel genes.

**S10 Table: Genes coding for histone methyltransferase in the mealybug genome.** *The pathways are those of Drosophila which are controlled by the histone methyltransferase. **The numbers refer to the number in the reference list given below.

S11 Table: Comparison of the number of chromatin remodeling genes in various insects.

**S12 Table: List of Chromatin Remodelers conserved in mealybugs and the potential histone modification they identify.**

**S13 Table: List of differentially expressed epigenetic modifiers between male and female mealybugs.** *A positive Log FC (fold change) value indicates higher expression in males while a negative Log FC value indicates higher expression in females.

**S14 Table: Expression of epigenetic modifier genes in male and female mealybugs in terms of average TPM values.** No differential expression observed. *A positive Log FC value indicates higher expression in males while a negative Log FC value indicates higher expression in females

S15 Table: Proteins involved in X inactivation that are shared between mammals and mealybugs (Mhir and Pcit).

**S16 Table: Primers (all written 5’ to 3’).** *, # and $ were used as primer sets for the long PCR sets represented as 1, 2 and 3 in Figure 2 (Validation of the Mhir assembly).

S17 Table: Complete Transcriptome Data for adult male and female *M. hirsutus*.

